# Decoding Multicellular Communication Motifs from Spatial Transcriptomics with ALARMIST

**DOI:** 10.64898/2026.05.21.726986

**Authors:** Jiayi Fan, John Hood, James Strong, Jeffrey F. Quinn, Yibo Dai, Data Science TeamLab, Aaron Schein, Kenny Kwok Hei Yu, Wesley Tansey

**Affiliations:** Computational Oncology, Memorial Sloan Kettering Cancer Center; Department of Statistics, University of Chicago; Department of Neurosurgery, Memorial Sloan Kettering Cancer Center; Department of Genomic Medicine, University of Texas MD Anderson Cancer Center; Break Through Cancer

## Abstract

Cellular organization is driven by recurrent, coordinated interactions between multiple cell types, each sending and receiving multiple signals. Existing computational methods for spatial profiling data consider only individual ligand–receptor interactions and fail to capture the higher-order interactions governing the tissue microenvironment. To address this gap, we developed ALARMIST (Assessment of Ligand And Receptor Motifs And Impacts in Spatial Transcriptomics), a probabilistic framework that infers interpretable multicellular communication patterns from spatial data. ALARMIST decomposes neighborhood-level signaling patterns into motifs: recurrent communication subnetworks involving multiple cell types and sets of enriched ligand-receptor interactions. For each cell, ALARMIST identifies its active motifs and estimates the downstream phenotypic effects of each motif on active cells. We applied alarmist to spatial datasets of lung adenocarcinoma (LUAD) and glioblastoma (GBM) to identify microenvironmental drivers of tumor progression. In paired LUAD and adenocarcinoma-in-situ (AIS) samples, ALARMIST identified an immune-active vascular motif at the tumor-normal boundary and implicated motif-active plasmacytoid dendritic cells as drivers of inflammation in early carcinogenesis. In matched low- and high-grade glioma samples, ALARMIST identified a hub-and-spoke motif centered on a malignant macrophage subpopulation, implicating a GRN-SORT1 signaling axis with a downstream impact gene set predictive of survival in low-grade glioma patients. Code for ALARMIST is available at https://github.com/tansey-lab/alarmist.

## Introduction

Tissue and organ function arise not from individual cells acting in isolation, but from the coordinated organization of diverse cell types. They collectively maintain homeostasis or, under disease conditions, become remodeled to drive pathological processes. For example, tumors induce an immunosuppressive microenvironment through a multi-signal program involving both direct signaling to effector cells via immune checkpoint receptors, as well as indirect suppressive signaling through macrophage polarization and fibroblast-induced remodeling of the extracellular matrix. ^1,2^ In kidney fibrosis, injured tubular epithelial cells secrete TGF-*β*, Wnt-signaling ligands, and inflammatory cytokines that activate fibroblasts and recruit immune cells, while fibroblasts signal through integrins, EGFR, and TLR4 on neighboring epithelial, fibroblast, and immune cells to reinforce a spatially confined fibrogenic niche. ^3^ Thus, even in highly dysfunctional tissue states, the pathology is still driven by higher-order, albeit malignant, communication programs driven in large part by sets of ligand-receptor interactions (LRIs) between multiple cell types.

A range of computational methods have been developed to infer LRIs from spatial transcriptomics data. Methods originally developed for scRNA-seq, including CellChat ^4^ and CellPhoneDB ^5^, have been adapted to spatial data by restricting analysis to nearby cell types. Spatially-aware methods have also been developed, including graph- and statistics-based approaches such as Giotto ^6^ and SpatialDM ^7^, optimal-transport frameworks such as COMMOT ^8^, and deep learning methods such as DeepTalk ^9^ and GITIII ^10^. Despite their methodological differences, these approaches largely score each LRI independently and return ranked lists of individually significant interactions, rather than identifying coordinated higher-order communication programs and recurring spatial signaling structure. Other methods have instead focused on representing cell-cell communication as multicellular networks. Connectome ^11,12^ represents cell-cell communication as cell-type-directed networks weighted by ligand-receptor co-expression. NICHES ^13^ reformulates this at single-cell resolution, treating each sender-receiver cell pair as an observation in a cell-pair by ligand-receptor matrix. In both cases, pattern discovery relies on graph centrality or post hoc clustering rather than explicit modeling of latent programs.

Factorization-based approaches have begun to address this limitation by extracting latent communication programs across samples. For example, Tensor-cell2cell ^14^, integrated with LIANA ^15^, identifies LRI patterns across samples, conditions, and cell-type pairs, but does not model spatial structure and aggregates expression at the cell-type level, losing single cell specificity. More recently, COMPOTES ^16^ introduced a spatially aware factorization framework that identifies recurrent multicellular programs across large patient cohorts, but it does not explicitly distinguish sender and receiver cell-type identities within each spatial unit. Across all methods, there is an inability to decompose LRIs at the single cell level, with cell type and spatial awareness, and in a manner that recovers the higher-order communication patterns that reflect the way multicellular biological systems function.

Here we present ALARMIST (Assessment of Ligand And Receptor Motifs and Impacts in Spatial Transcriptomics), a computational framework that identifies coordinated cell–cell communication programs from spatial transcriptomics data. ALARMIST jointly models all LRIs across tissue space using Bayesian Poisson tensor factorization to discover communication motifs: recurring sets of cell types and LRIs that co-activate in the same spatial microenvironment. Each motif captures not only which ligand-receptor pairs are involved but also the specific sender–receiver cell-type combinations that participate. Motifs are projected to single-cell resolution and linked to changes in expression of non-ligand/receptor genes, enabling interpretation of how local signaling environments shape cells’ transcriptional states. Using matched Xenium and CosMx profiling of consecutive tissue sections from three cancer types, we show that ALARMIST motifs, their spatial organization, and their downstream transcriptional effects are concordant across platforms, with residual discordance attributable to platform-level differences in cell-type gene detection. On Xenium 5K datasets from lung adenocarcinoma and glioblastoma, ALARMIST uncovered motifs driving disease progression through reshaping of the tumor microenvironment. In lung adenocarcinoma, it resolved a cancer precursor vascular program and implicated plasmacytoid dendritic cells as drivers of local inflammation. In glioma, it identified a malignancy associated glioma macrophage-centered motif engaged in a feedback loop with an aggressive tumor subtype, and a downstream impact gene set predictive of survival in low grade glioma patients.

## Results

### Overview of ALARMIST

Cell-cell communication in the tissue microenvironment is inherently multicellular. Multiple cell types signal to one another simultaneously, forming coordinated interaction patterns that span the local tissue architecture. Further, cells can be participating in multiple coordinated activities, such as ECM remodeling, vascular stabilization, and antigen presentation, simultaneously. The resulting spatial snapshot observed in a spatial transcriptomics dataset is then a complex admixture of signaling patterns (Figure 1a). Standard LRI analysis methods reduce this complexity to isolated pairwise interactions, discarding local spatial context and thus lacking the higher-order organization of communication (Figure 1b). ALARMIST addresses this by discovering multicellular LRI motifs capturing recurring, spatially coherent units of coordinated signaling among multiple cell types that constitute the building blocks of tissue communication (Figure 1c). ALARMIST also assigns motif activities to individual cells, revealing spatial structures in the tissue that are organized by cell-cell communication (Figure 1d).

**Figure 1.**
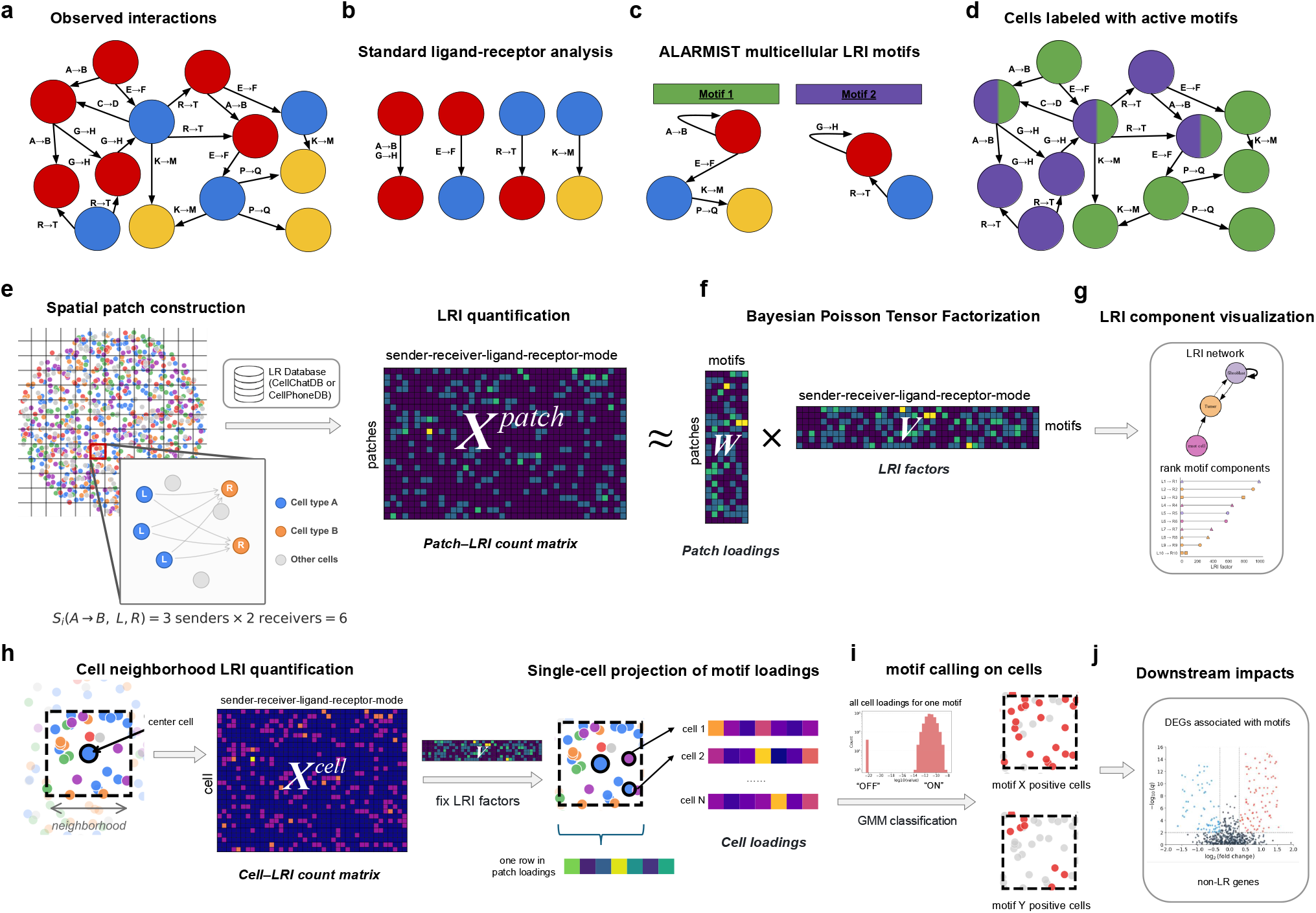
Overview of the ALARMIST framework.**a.**Cell-cell communication in the tissue microenvironment involves multiple cell types signaling simultaneously, forming coordinated interaction patterns across the local tissue architecture. **b**. Standard LRI analysis reduces this complexity to isolated pairwise interactions, discarding spatial context. **c**. ALARMIST discovers multicellular LRI motifs: recurring, spatially coherent units of coordinated signaling among multiple cell types and LR pairs. **d**. Motif activities are assigned to individual cells, revealing spatial structures organized by cell-cell communication. **e**. Spatial patch construction and LRI quantification. Tissue is discretized into spatial patches, and LRI counts are computed for each patch using curated databases (CellChatDB or CellPhoneDB). For a given patch, the count for each sender-receiver cell type pair and ligand-receptor pair is defined as the product of ligand-expressing sender cells and receptor-expressing receiver cells. **f**. Bayesian Poisson matrix factorization (BPTF) decomposes the patch-LRI count matrix (*X*^*patch*^) into patch loadings (*W*) and LRI factors (*V*), identifying *K* latent communication motifs. Spatial permutation tests assess statistical significance. **g**. Motif LRI component visualization. Each motif’s LRI composition is visualized as a cell type interaction network and ranked by component contribution. **h**. Cell neighborhood LRI quantification and single-cell projection. A cell-LRI count matrix (*X*^*cell*^) is constructed from each cell’s local neighborhood. Motif loadings are projected to single-cell resolution by refitting the model with LRI factors (*V*) fixed from the patch-level decomposition. **i**. Motif activation calling. Per-cell loadings for each motif are classified into discrete ON/OFF states via Gaussian mixture model thresholding, yielding spatial maps of motif-positive cells. **j**. Downstream impact analysis. Poisson generalized linear models identify genes whose expression is associated with each motif, revealing downstream transcriptional programs beyond ligand-receptor genes.

ALARMIST formulates cell-cell communication motif discovery as a matrix factorization problem. To construct the matrix, LRI events are quantified within spatial local neighborhoods (Figure 1e). The tissue is divided into small patches by overlaying a regular grid, so each patch defines a spatial neighborhood. For each patch, LRI counts are quantified by counting co-expression events between ligand-expressing cells (senders) and receptor-expressing cells (receivers) across all cell type pairs. Specifically, the LRI count *S*_*i*_(*A, B, L, R*) for patch *i* is defined as the number of sender cells multiplied by the number of receiver cells. For example, if a patch contains 3 cells of type *A* expressing ligand *L* and 2 cells of type *B* expressing receptor *R*, the interaction count is *S* = 3 × 2 = 6. LRI definitions are drawn from curated databases such as CellChatDB^4^ or CellPhoneDB ^17^. This produces a patch-by-LRI count matrix *X* that is typically large; in our datasets, *X* is on the order of 100K × 100K.

The size of the patch-LRI matrix presents a computational challenge, both in terms of storing the matrix and factorizing it. Fortunately, the matrix is also generally extremely sparse. ALARMIST leverages this sparsity through the use of Bayesian Poisson tensor factorization (BPTF)^18^, which scales computationally with the number of nonzero entries. ALARMIST uses BPTF to decompose this count matrix into a small number of latent communication motifs (Figure 1f). The factorization yields two matrices: a patch loading matrix *W*, indicating how strongly each patch *i* expresses motif *k*; and an LRI factor matrix *V*, capturing which LRIs and cell type pairs define motif *k*. The Poisson likelihood is a natural choice for modeling LRI counts as they are non-negative integers. The BPTF Poisson–Gamma construction provides shrinkage that is well-suited to our high-dimensional, extremely sparse count matrix, improving robustness compared to non-Bayesian low-rank decompositions. The motifs are visualized as ranked lists of top LRIs and network plots of cell types constructed from summing the top LRI factors (Figure 1g).

Motifs discovered at the patch level are projected to single-cell resolution (Figure 1h). Because patch-level loadings aggregate over all cells within a patch, we refine motif activities to individual cells by constructing a cell-centered neighborhood for each cell and quantifying its LRI counts in the same manner as above, producing a cell-by-LRI matrix *X*_*cell*_. Per-cell motif loadings are then inferred by fixing the LRI factor matrix *V*, learned at the patch level, and fitting only the cell loading parameters *U*. This projection shows motif activities that reflect individual cell participation rather than patch-level averages. The resulting cell-level loadings follow a bimodal distribution on the log scale, with one mode near zero and another at higher values, consistent with a mixture of inactive and active cells (Figure 1i). Accordingly, a Gaussian mixture model is fitted to classify cell loadings into discrete motif activation states (ON or OFF) for each motif.

After assigning motifs to cells, ALARMIST conducts a downstream analysis to link motif activity to transcriptional changes (Figure 1j). Poisson generalized linear models are used to identify genes (excluding the ligand and receptor genes already used in motif construction) whose expression is significantly associated with each motif. To avoid confounding from transcript diffusion across nearby cell types, marker genes of other cell types are optionally excluded from visualization in volcano and forest plots.

ALARMIST leaves the patch size and number of latent motifs as user-specified hyperparameters. As a default, patch sizes are set to 50 microns to capture both juxtacrine and paracrine signaling. In sparse tissues, such as the lung tissue in Fig. 4, we recommend increasing the patch size to increase the number of cells in each patch and reduce variance. The choice of number of motifs is analogous to the number of latent factors in non-negative matrix factorization. The choice is generally robust but should be tuned by the user to balance complexity of the programs with adequate coverage of the biological processes at work in each tissue.

### ALARMIST motifs capture higher-order communication patterns with single-cell specificity

We first validated ALARMIST in a simulation where we constructed synthetic motifs from scRNA-seq reference data ^19^(Fig. 2a). From the original data, *Y* ^ref^, consisting of 8,288 cells, we constructed ten synthetic motifs, each with three to five distinct cell types, out of 16 annotated types in the scRNA-seq dataset (Supplementary Figure 2). Each motif was defined by a directed network between cell types (Fig. 2b). For each edge (from sender to receiver cell type), we selected five associated LRIs using CellPhoneDB with a bias towards higher expression ligands and receptors in senders and receivers, respectively (Methods). For each participating cell type, we also selected ten non-ligand/receptor marker genes to be differentially expressed in an active versus inactive motif. For each differentially expressed gene set, five were assigned to be upregulated and five were downregulated. To minimize impact on the overall gene expression distribution of each cell type, we used biased permutations of the observed gene expression values to impose differential expression and LRI activity in motifs. Motif active regions were sampled from a Gaussian process to ensure spatially contiguous regions of activity to better reflect tissue structure (Supplementary Figure 1b). A motif is defined by the spatial regions where it is active, the cell type interaction network, LRI set, and the differentially expressed genes. We evaluated ALARMIST’s ability to recover each element of the motifs as well as the spatial motif structure. We first assessed whether ALARMIST could recover the directed cell type networks defining each motif. We benchmarked ALARMIST against three recent methods, COMPOTES ^16^, Tensor-cell2cell ^14^, and NICHES ^13^, comparing each method’s estimated networks to the ground truth using cosine similarity (Methods). ALARMIST consistently recovered the ground-truth interaction networks, achieving a mean cosine similarity of approximately 0.8 across dataset sizes, whereas NICHES, Tensor-cell2cell and COMPOTES fell below 0.6, 0.5, and 0.4, respectively (Fig. 2c,d). The top-ranked LRIs for each recovered motif largely matched the ground-truth active interactions (Fig. 2i), confirming that ALARMIST correctly identified both the cellular participants and the molecular channels of each program.

**Figure 2:**
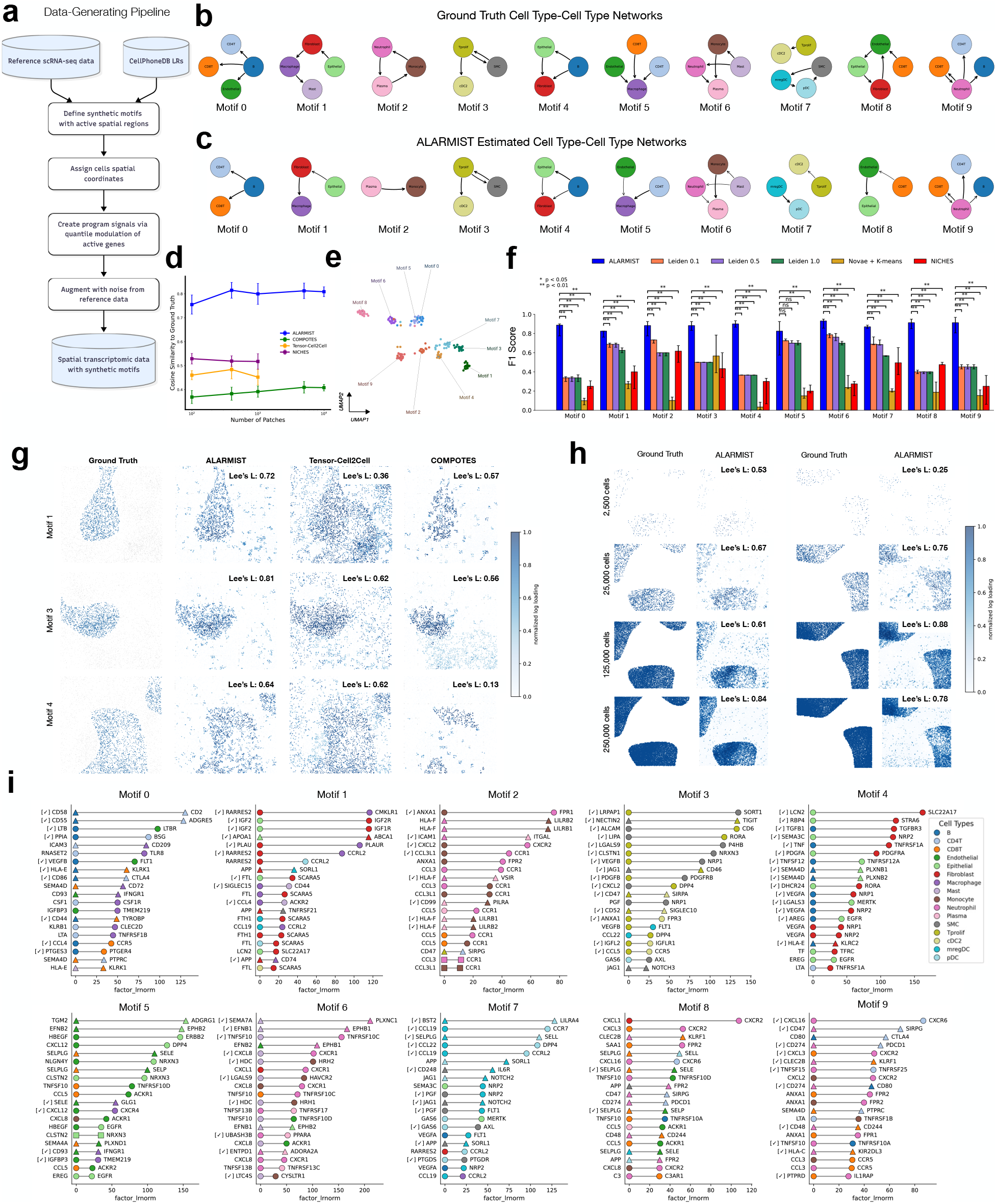
ALARMIST recovers ground-truth spatial signaling motifs in synthetic benchmarks. **a**. Schematic of the synthetic data-generating pipeline. **b**. Ground-truth cell-type interaction networks for the ten synthetic motifs. **c**. ALARMIST-estimated cell-type interaction networks for the same ten motifs. Edges weighted by normalized LRI motif loadings summed across LRIs. **d**. Cosine similarity between recovered and ground-truth LRI factor vectors as a function of the number of spatial patches, for ALARMIST, Tensor-Cell2Cell, COMPOTES, and NICHES. Points show the median and error bars show the interquartile range across random initializations. **e**. UMAP of motif LRI factor vectors across all methods and replicates, colored by ground-truth motif identity. **f**. Per-motif F1 score for cell-level motif assignment, comparing ALARMIST against Leiden clustering at three resolutions (0.1, 0.5, 1.0), Novae + *K*-means, and NICHES. Bars show the median and error bars show the interquartile range across random initializations. *P* -values from two-sided Wilcoxon signed-rank tests (^*^*P* < 0.05; ^**^*P* < 0.01; ns, not significant). **g**. Spatial structure recovered by each method. ALARMIST yields cell loadings, while Tensor-Cell2Cell and COMPOTES yield patch level-loadings. Color indicates normalized log loading. Lee’s *L* between the recovered and ground-truth spatial pattern is shown in each panel. **h**. Recovery performance at varying cell counts, shown for two representative motifs. Lee’s *L* between ALARMIST and ground truth is shown in each panel. **i**. Top LRIs of each ALARMIST-recovered motif ranked by motif loading. Markers indicate sender (left) and receiver (right) cell types; marker color denotes cell type, and marker shape denotes signaling type (square, autocrine; circle, paracrine; triangle, juxtacrine). A [✓] prefix indicates the LRI was present in the corresponding ground-truth motif.

In real tissues, the same cell type often participates in multiple communication programs through different sets of ligand-receptor interactions. Therefore it is important to resolve program-specific molecular signatures despite shared cellular participants. To investigate this, we examined ALARMIST’s LRI embedding space, where each LRI receives a K-dimensional loading vector across the inferred motifs (number of motifs, K=20; Fig. 2e). UMAP visualization of the log-transformed embeddings revealed that active LRIs clustered into visually distinct groups corresponding to their ground-truth motif assignments for nearly all ten motifs, indicating that ALARMIST correctly attributed each LRI to its originating program even when motifs shared cell types. Motifs 0 and 5 showed partial entanglement, likely reflecting ALARMIST’s omission of CD8T from its estimated network for motif 0 (Fig. 2c). The full embedding space, including inactive interactions, is shown in Supplementary Figure 1a.

We next assessed each motif’s differentially expressed gene recovery using the ALARMIST downstream impacts. We compared the ALARMIST results to three different parameter choices for Leiden clustering and K-means clustered embeddings from Novae, ^20^ a spatial foundation model (Methods). These baselines assign each cell to a single cluster, while ALARMIST provides continuous cell loadings that allow a cell to participate in multiple motifs simultaneously. For each method and motif, we selected the top ten non-LRI genes by absolute log-fold change and compared them to the ten ground-truth marker genes. ALARMIST achieved the highest F1 score across all ten motifs (Fig. 2f). For motif 5, where network recovery had also been incomplete (Fig. 2c), ALARMIST did not significantly outperform the Leiden baselines. Overall, the ALARMIST F1 scores were over 0.6 for every motif, whereas every other baseline had worst-case performance below 0.35.

Finally, we evaluated ALARMIST’s ability to recover the spatial structure underlying each motif. A key distinction is that ALARMIST assigns loadings at single-cell resolution, whereas Tensor-cell2cell and COMPOTES merely operate at the patch level. We quantified spatial recovery using Lee’s L ^21^, a bivariate spatial correlation coefficient, between the ground-truth cell states and each method’s estimated loadings. ALARMIST consistently achieved the highest correlation values across motifs (Fig. 2g, Supplementary Figure 1d), while COMPOTES and Tensor-cell2cell showed comparatively poor recovery for specific motifs (e.g., motifs 4 and 1, respectively). We also evaluated the robustness of the ALARMIST spatial recovery to varying cell densities (Figure 2h). Across varying cell counts from 2.5K to 250K we found that cell-level loadings faithfully recapitulated the ground-truth active regions even at lower cell densities (Supplementary Figure 1c).

### Cross-platform concordance of ALARMIST motifs and spatial niches

To evaluate the robustness of ALARMIST across spatial transcriptomics platforms, we applied it to matched Xenium 5K and CosMx 6K datasets from three cancer types (COAD, OV, HCC), where consecutive tissue sections from the same specimen were profiled on each platform ^19^ (Figure 3a). All six samples were processed with identical parameters (50 *µ*m patches, *K* = 20 motifs).

**Figure 3:**
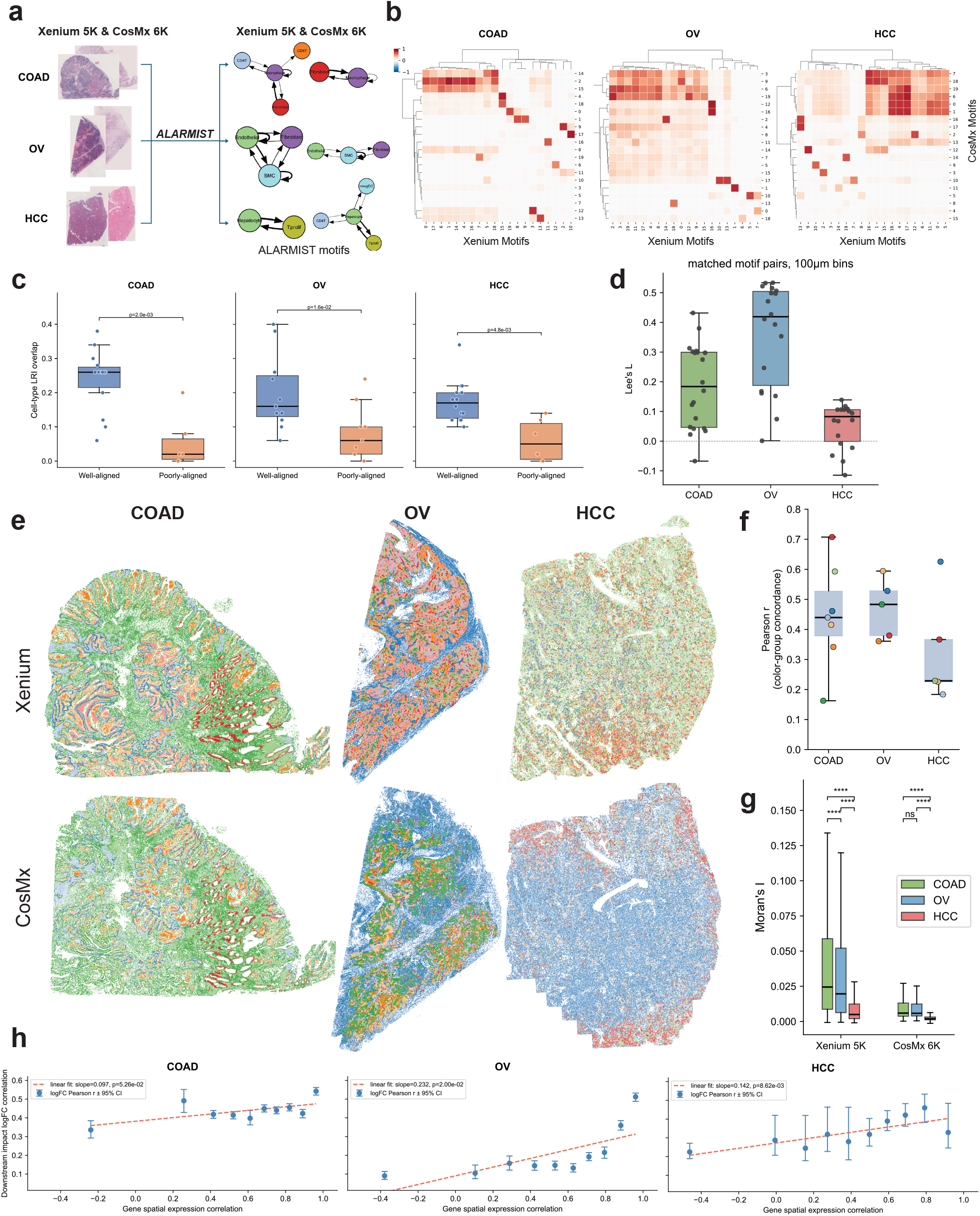
Cross-platform concordance of ALARMIST motifs and spatial niches. **a**. Schematic of the cross-platform comparison. **b**. Heatmaps of pairwise cosine similarity between Xenium and CosMx motif LRI factor vectors for each cancer type. Color scale indicates cosine similarity. **c**. Cell-type-LRI overlap rate between the effective LRI pool of Xenium motifs and the top 50 cell-type-LRIs of each CosMx motif, stratified by alignment status (well-aligned versus poorly aligned). *p*-values from two-sided Mann-Whitney *U* tests. **d**. Lee’s *L* statistic for matched motif pairs, computed on the proportion of motif-positive cells per 100 *µ*m bin across the platform intersection using queen contiguity spatial weights. Each dot represents one matched motif pair. **e**. Spatial maps of niche clusters for Xenium and CosMx within the platform overlap region. CosMx niches are colored by their most correlated Xenium niche. **f**. Pearson correlation of niche proportion vectors across 500 *µ*m grid squares between the two platforms. Each dot represents one matched niche pair, colored as in (e). **g**. Moran’s *I* of genes shared between platforms, computed with 20 nearest neighbors. Each dot represents one gene. *P* -values from two-sided Mann-Whitney *U* tests. **h**. Relationship between gene expression concordance and downstream impact concordance across platforms. Genes were binned into 10 quantiles by spatial expression correlation (*x* axis). *y* axis shows the Pearson correlation of matched-motif log-fold changes within each bin. Error bars indicate 95% bootstrap confidence intervals. Line shows linear regression fit with slope and *p*-value.

We first compared the LRI composition of motifs across platforms by computing cosine similarity between each pair of Xenium and CosMx motif LRI factor vectors, restricted to cell-type-LRI combinations shared between platforms. Many motif pairs showed high concordance, with well-matched pairs reaching cosine similarities of approximately 0.7 (Figure 3b, Supplementary Figure 3). Using a threshold of 0.4, we classified each motif as well-aligned the best-matched motif on the opposing platform exceeded this threshold, and poorly aligned otherwise. Most Xenium motifs had a CosMx counterpart; in contrast, 5, 8, and 6 CosMx motifs were poorly aligned in COAD, OV, and HCC, respectively.

To understand this asymmetry, we examined whether the top 50 enriched LRIs of poorly aligned CosMx motifs overlapped with the pooled effective LRIs of Xenium motifs. We considered three different kinds of overlap: at the level of individual ligand or receptor genes, ligand-receptor pairs, and cell-type-LRI combinations. Gene-level and LR-pair-level overlap showed no significant difference between well-aligned and poorly aligned motifs (Supplementary Figure 3). The difference emerged only at the cell-type-LRI level (Mann-Whitney *U* test; Figure 3c), suggesting that platform discordance between ALARMIST motifs arises from differences in which cell types express particular LRIs.

We next asked whether matched motifs exhibited concordant spatial distributions. After aligning the two platforms via a similarity transformation (Supplementary Figure 4a), we computed Lee’s *L* statistic for each matched motif pair on the proportion of motif-positive cells per 100 *µ*m bin across the platform intersections (Figure 3d). Spatial concordance was moderate overall, with OV showing the strongest agreement, followed by COAD and HCC. Notably, Lee’s *L* values for matched motif pairs exceeded those for same-cell-type comparisons across platforms (Supplementary Figure 4b), implying that ALARMIST motifs capture spatially coherent signals beyond cell-type composition alone.

To evaluate whether motif-defined spatial niches were concordant, we clustered cells by their normalized loadings across matched motifs using *K*-means. The choice of *K* was determined by elbow analysis on Xenium and applied to both platforms (Supplementary Figure 4c). Niche proportions were computed in 500 *µ*m grid squares across the platform overlap, and each CosMx niche was mapped to its most correlated Xenium niche by Pearson correlation of proportion vectors (Figure 3e, Supplementary Figure 4d). COAD and OV showed the strongest niche concordance, whereas HCC was more divergent (Figure 3f, Supplementary Figure 4e). This distinction paralleled the spatial structure of the underlying gene expression: Moran’s *I* of shared genes was significantly lower in HCC than in the other two cancers (Mann-Whitney *U* test; Figure 3g), consistent with reduced tissue-level spatial organization.

Finally, we assessed whether matched motifs yielded consistent downstream transcriptional impacts. For each shared gene, we computed the Pearson correlation of pseudo-bulk expression between platforms. For each matched motif pair and shared cell type, we computed the correlation of log-fold differential expression changes between platforms for genes significant on either platform. Binning genes by their expression correlation revealed a positive relationship: genes with more concordant spatial expression showed more concordant downstream impact estimates (Figure 3h,). This held for OV and HCC (linear regression Pearson *r* < 0.05), while COAD showed uniformly high impact concordance across all bins with a less pronounced positive slope (Pearson *r* = 0.0526).

Overall, between both platforms we found ALARMIST motifs, spatial localization, and downstream impacts to be robust to moderate technical differences. Where ALARMIST results diverged most were attributable to either lack of substantial overlap in the underlying panels and cell typing, or due to relatively less rich tissue structure in which to detect spatially localized patterns.

### ALARMIST reveals divergent vascular microenvironments underlying immune remodeling in lung adenocarcinoma progression

We applied ALARMIST to study cell-cell communication patterns during lung adenocarcinoma (LUAD) progression. We analyzed Xenium 5k data of four samples from two patients, each with matched precursor adenocarcinoma-in-situ (AIS) and invasive LUAD tissue sections ^22^ (Supplementary Figure 8a,b). The dataset comprised over 1.6 million cells profiled with a 5096-gene panel, yielding 772 detectable LRI pairs from CellChatDB v2.0 and enabling analysis of LRI organization across the pre-invasive to invasive transition.

We used ALARMIST to identify 25 LRI motifs across the dataset (Supplementary Figure 5). We noted two motifs capturing contrasting vascular microenvironments: a healthy vasculature motif and a tumor vasculature motif (Figure 4b,f). The healthy vasculature motif mainly involves alveolar epithelial cells, fibroblasts, and vascular endothelium (Figure 4b), whereas the tumor vasculature motif is dominated by tumor epithelium, pericytes, and vascular endothelium (Figure 4f). Given the lack of tumor signaling in the healthy state and centrality of tumor cells in the other motif, we hypothesized that the two different motifs may capture spatiotemporal differences in disease progression.

**Figure 4:**
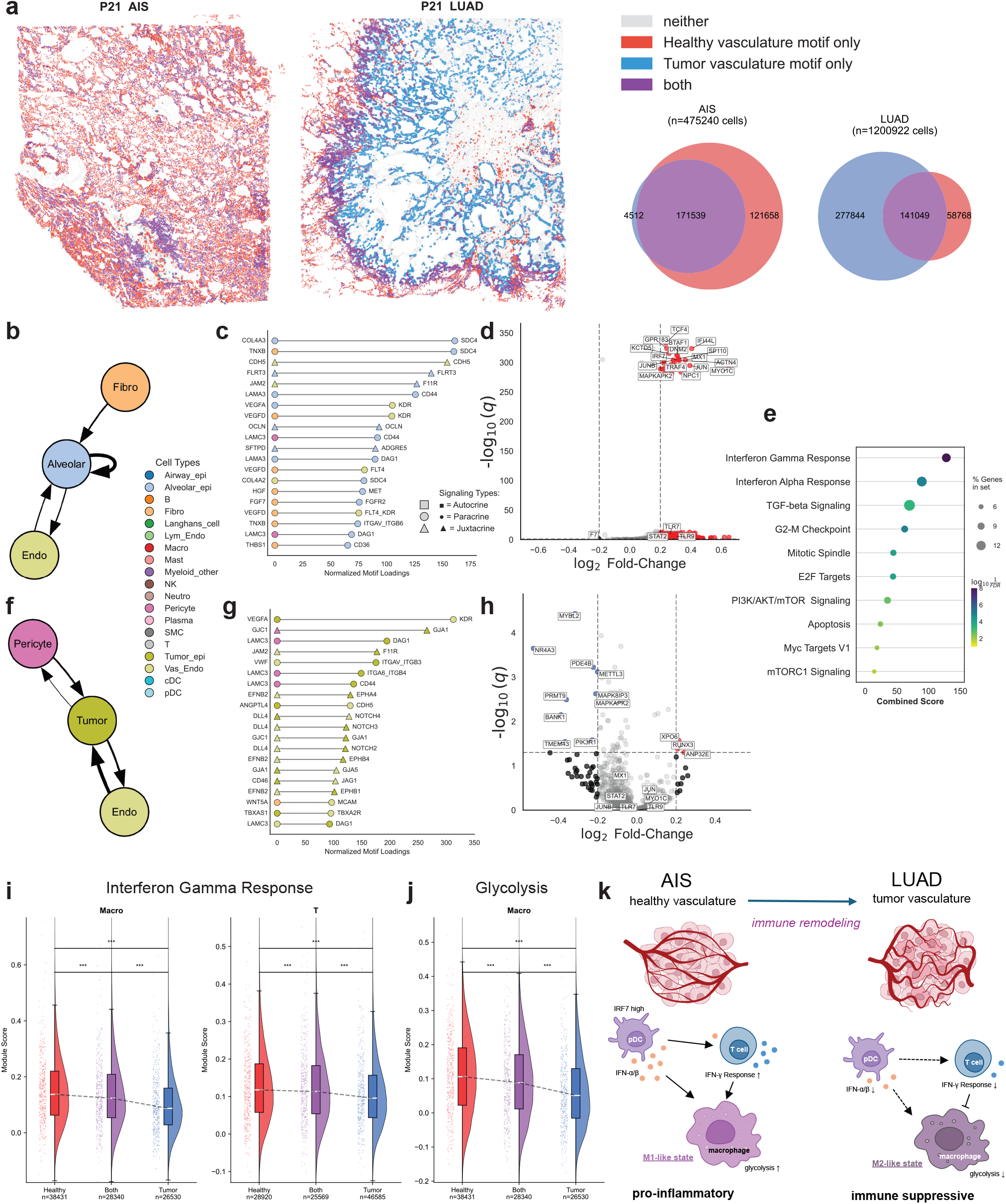
Vascular and immune remodeling during progression from AIS to LUAD. **a.** Representative P21 AIS and LUAD sections showing cells positive for the healthy vasculature motif, the tumor vasculature motif, or both. Venn diagrams summarize the per-section cell composition across the three states. **b**. Cell-type interaction network of the healthy vasculature motif. Edges weighted by normalized LRI motif loadings summed across LRIs. **c**. Top 20 ligand-receptor interactions (LRIs) of the healthy vasculature motif, ranked by LRI factor value. **d**. Downstream impact of healthy vasculature motif on pDCs. Each dot represents one gene; effect sizes (log_2_FC) are *β* coefficients from a per-gene Poisson GLM with motif state as the covariate, and *q*-values are from Wald tests adjusted using Benjamini-Hochberg. Significance thresholds: |log_2_ FC| > 0.2, *q* < 0.05. **e**. Gene set enrichment analysis (GSEA) of pDC downstream impacts from (d) against MSigDB Hallmark gene sets, highlighting interferon-related programs. **f**. Cell-type interaction network of the tumor vasculature motif. Edges weighted as in (b). **g**. Top 20 LRIs of the tumor vasculature motif, ranked as in (c). **h**. Downstream impact of tumor vasculature motif on pDCs, computed as in (d). **i**,**j**. Hallmark Interferon-*γ* response (i) and glycolysis (j) module scores in macrophages and T cells stratified by motif state. Box plots show median (center line), interquartile range (box), and 1.5 × IQR (whiskers). *p*-values from two-sided Mann-Whitney *U* tests. **k**. Model summarizing the transition from a pro-inflammatory, interferon-active vascular niche in AIS to an immunosuppressive tumor vasculature niche in LUAD.

Spatial mapping of motif states in cells revealed different distribution patterns across disease stages (Figure 4a, Supplementary Figure 8b,c). In AIS tissue, the healthy vasculature motif was broadly expressed, while tumor regions were characterized by co-expression of both motifs (65–69% in tumor region) rather than a switch to the tumor vasculature motif alone (Supplementary Figure 9). In LUAD tissue, the tumor vasculature motif dominated the tumor core, while the remaining healthy vasculature motif was largely confined to normal tissue. Co-expression of both motifs was enriched at the tumor-normal interface. These patterns suggested that progression from AIS to LUAD is accompanied by an erosion of healthy vasculature signaling programs.

We next reasoned that the vascularization motifs may reflect different microenvironmental drivers of tumorigenesis and invasion. To investigate this, we first examined the LRI content of each vascular motif (Figure 4c,g). In the healthy vasculature motif, top-ranked interactions include fibroblast-derived epithelial support signals (*HGF* → *MET, FGF7* → *FGFR2, FGF2* → *FGFR2*) and extracellular matrix interactions (*COL4A3* → *SDC4, THBS1* → *SDC4*). These interactions are consistent with known roles of mesenchymal-epithelial crosstalk in maintaining alveolar homeostasis. In the tumor vasculature motif, pro-angiogenic signaling dominates: *VEGFA* → *KDR* is the top-ranked interaction, accompanied by Notch pathway components (*DLL4* → *NOTCH1, JAG1* → *NOTCH2*) and additional angiogenic signals (*ANGPTL4* → *CDH5*). Notably, fibroblast-derived support signals are absent, indicating disruption of the homeostatic niche.

We next assessed the ALARMIST downstream impact estimates on each cell type in the microenvironment. Both motifs produced significant differentially expressed genes in all 19 cell types tested but showed inverted cellular polarity. The tumor vasculature motif upregulated tumor and stromal compartments (tumor epithelium, pericytes, vascular endothelium) while predominantly suppressing immune compartments (Supplementary Figure 11). The healthy vasculature motif showed the opposite: it broadly upregulated immune compartments (T, B, NK, dendritic, plasma, and macrophages) and alveolar epithelium, while uniquely suppressing tumor epithelium (Supplementary Figure 12). Intriguingly, we noted that conventional dendritic cells (cDCs) had no detectable downstream impact genes, whereas plasmacytoid dendritic cells (pDCs) had a number of positively- and negatively-impacted genes in both motifs (Figure 4d,h). In healthy vasculature regions, pDCs showed significant upregulation of *IRF7* (log_2_ FC = 0.317, *q* < 10^−300^), the master regulator of type I interferon (IFN) production, along with *TLR7* (log_2_ FC = 0.261, *q* = 1.8 × 10^−7^) and *TLR9* (log_2_ FC = 0.297, *q* = 1.5 × 10^−6^), the endosomal sensors that initiate IFN responses to nucleic acids. Interferon-stimulated genes including *MX1, OAS1*, and *STAT2* were also elevated. Gene set enrichment analysis confirmed significant enrichment of Hallmark Interferon Alpha Response in healthy vasculature pDCs (Figure 4e). In contrast, pDCs in tumor vasculature regions showed downregulation of *IRF7* (log_2_ FC = −0.082, *q* = 2 × 10^−4^) and *METTL3* (log_2_ FC = −0.201, *q* = 7.6 × 10^−4^), the m6A RNA methyltransferase required for dendritic cell activation, with no significant changes for interferon-stimulated genes.

Based on the ALARMIST motifs and impacts, we hypothesized that pDCs in AIS act as inflammatory drivers in early lung cancer. To further investigate this, we examined whether the inflammatory pDC states were concordant with differences in effector immune function between motifs (Supplementary Figure 13). IFN-*γ* response module scores were significantly higher in T cells and macrophages located in healthy vasculature regions compared to tumor vasculature regions (Mann-Whitney U test; *p* < 0.001; Figure 4i), with cells in regions harboring both motifs showing intermediate scores but still significantly different from either side. Macrophages in healthy vasculature regions also displayed elevated glycolytic activity (Figure 4j), a metabolic hallmark of pro-inflammatory activation, whereas tumor vasculature macrophages showed reduced glycolysis. These patterns suggest coordinated immune activation in the healthy niche and immune suppression in the tumor niche, with an intermediate transitional state in regions of motif co-activation.

These results align with the original analysis, ^22^ which found an inflammatory niche in early lung cancer. The ALARMIST impacts, however, extended beyond these findings to implicate pDCs as potential drivers of tumor progression through IFN-stimulated inflammation in AIS (Figure 4k). Taken together, the ALARMIST results revealed that healthy vasculature in AIS enters an inflamed state where inflammatory pDCs in the fibroblast-epithelial-endothelial niche upregulate IFN and contribute to IFN-stimulated T cells and M1-like macrophages. In tumor vasculature, the loss of homeostatic support signals coincides with pDC functional impairment and dampened effector responses, establishing an immunosuppressive microenvironment.

The results also broadly match the line work showing chronic interferon stimulation acts as an escape mechanism for cancers treated with immune checkpoint blockade (ICB). ^23,24^ In particular, recent results show one ICB resistance mechanism is driven by reactivation of endogenous retroviral genes which lead to chronic interferon stimulation. ^25^ The canonical role of plasmacytoid dendritic cells is in viral response and pDCs are likely a contributor of interferon stimulation in the ICB setting. Building on this, the ALARMIST results suggests a provocative, biologically-rational hypothesis: chronic interferon may not only be a way to escape the immune system under ICB, but also a way to escape initial immune surveillance, thereby enabling tumor establishment.

### ALARMIST identifies an mGAM-centered spatial interaction motif enriched in transforming low grade glioma

Low grade gliomas (LGG) are a subset of primary brain cancers which harbor mutations in the gene for isocitrate dehydrogenase (IDH). LGGs are characterized by a diffuse, indolent yet progressive disease process with a propensity for high grade transformation. Once transformed, the the disease entity resembles glioblastoma (GBM), with a correspondingly poor prognosis. In contrast to GBM, which tends to occur in older adults, LGG typically occurs in adolescents and young adults. The drivers of LGG transformation remain unknown. Genomic differences between primary and recurrent samples do not adequately explain the transformation process. This has led to increased scrutiny of microenvironmental factors which may play a role in transformation.

Recent work ^26^ has identified a pathogenic macrophage subpopulation, termed malignancy-associated glioma macrophages (mGAMs), defined by a FOSL2-centered gene regulatory network and marked by surface co-expression of ANXA1 and HMOX1. This mGAM subpopulation was shown to be enriched in high-grade and actively transforming regions in LGG and co-culture of mGAMs with low grade glioma cells induces transition to a more aggressive, mesenchymal-like state.

The mesenchymal-like transcriptional state in gliomas is known to be associated with worse prognosis and an immunosuppressive, macrophage-rich microenvironment^27^, suggesting a causal role for mGAMs in tumor transformation and progression. The selective enrichment of mGAMs within high grade disease suggests that mGAM-tumor interactions may play an important role in disease progression, however how mGAMs coordinate with neighboring tumor cells through ligand–receptor interactions and what transcriptional consequences these interactions impose remains unknown.

To delineate the cellular crosstalk behavior connecting mGAMs and MES-like transformation in LGG, we profiled tissue microarray (TMA) samples from a select cohort of patients of five IDH-mutant LGG patients with tumors with evidence of active high grade transformation. Hematoxylin and eosin staining samples from this cohort of patients were retrieved and regions of low and high grade tumor were reviewed and annotated by a board certified neuropathologist (TB) before selectively sampled to create the TMA. Each patient therefore had paired low grade and high grade regions available for analysis (Figure 5a, Supplementary Figure 14). Tissues were spatially profiled using Xenium 5K. Cell types were annotated using a CellTypist model trained on the GBMap core atlas at annotation level 2 (^28,29^), identifying major malignant, immune, and vascular cell populations across samples (Figure 5b). Tumor cell subtypes were further annotated by computing module scores based on previously published marker gene signatures for the four canonical transcriptional states: mesenchymal-like, astrocyte-like, oligodendrocyte progenitor-like, and neural progenitor-like (MES-like, AC-like, OPC-like, NPC-like, respectively) ^30^. Macrophage were divided into mGAM and non-mGAM subpopulations using a FOSL2-centered module score. ^26^ In total, the dataset contained 100,197 cells profiled with the Xenium 5k panel (5,119 genes). Comparison with CellChatDB v2 identified 748 candidate LRIs.

**Figure 5:**
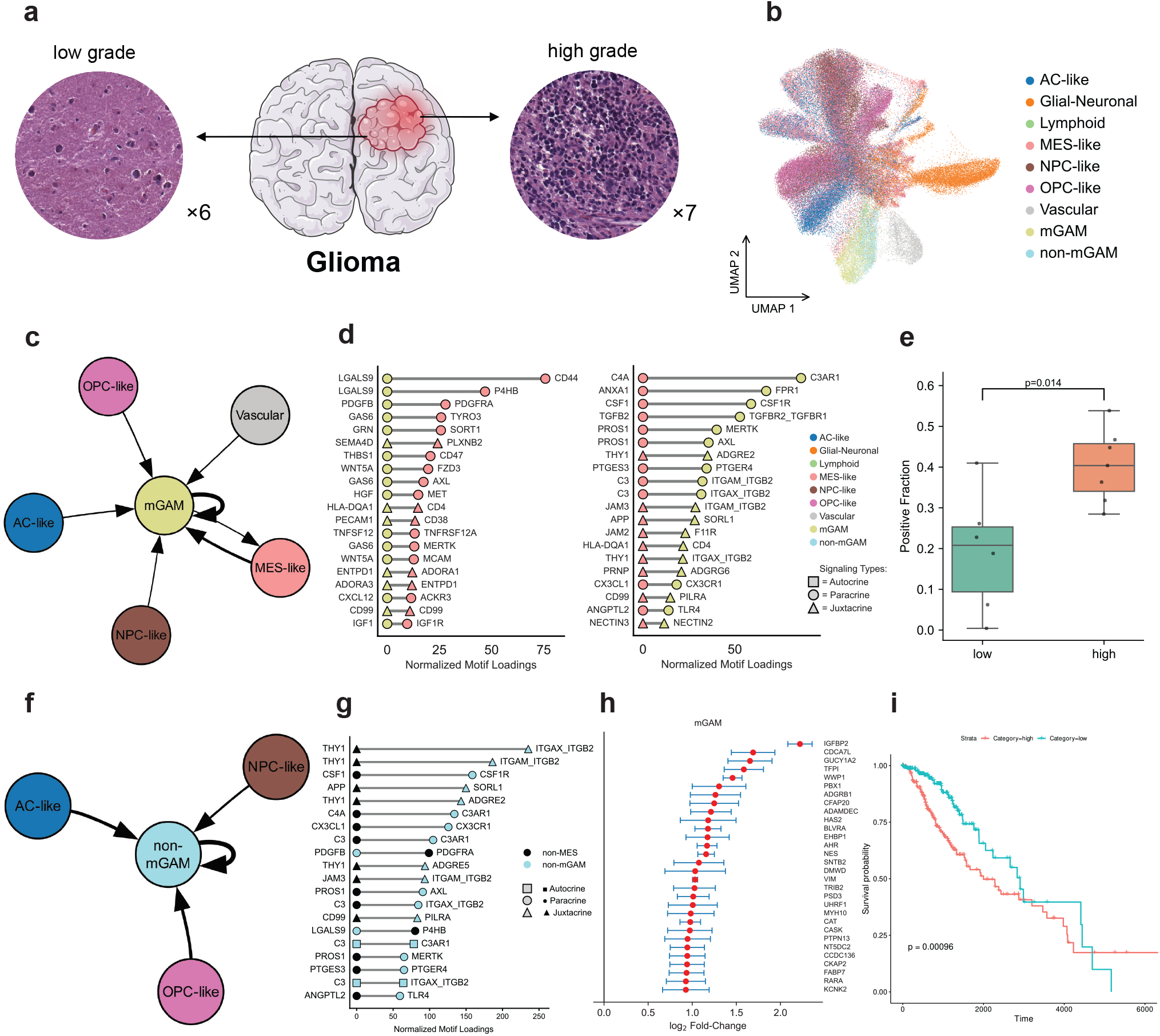
ALARMIST identifies an mGAM-centered spatial interaction motif enriched in transforming low grade glioma. **a**. Schematic of Xenium 5K profiling of glioma TMA samples (*n*=6 low-grade, *n*=7 high-grade cores) with representative H&E images. **b**. UMAP of 100,197 annotated cells. **c**. Cell type interaction network of the mGAM motif. Edges weighted by normalized LRI motif loadings summed across LRIs. **d**. Top 20 LRIs of the mGAM motif, split by direction: mGAM→MES-like (left) and MES-like→mGAM (right). **e**. Fraction of mGAM motif-positive cells in low-grade and high-grade samples. Box plots show median, IQR, and 1.5×IQR whiskers. *p*-value from two-sided Mann-Whitney *U* test. **f**,**g**. Cell type interaction network and top LRI components of a non-mGAM motif, which preferentially involves non-MES tumor states. **f**. Cell-type interaction network of the non-mGAM motif, weighted as in (c). **g**. Top 20 LRIs of the non-mGAM motif, as in (d). Non-MES-like tumor states are collapsed into a single non-MES category. **h**. Downstream impact of mGAM motif on mGAMs, showing top 30 gens ranked by log_2_ fold-change. Effect sizes are *β* coefficients from Poisson GLMs; error bars show 95% CI. All genes pass *q* < 0.001 (Wald test, Benjamini-Hochberg adjusted). **i**. Kaplan-Meier survival of TCGA LGG patients stratified by expression of the gene program in (h). *p*-value from log-rank test.

We applied ALARMIST, using 20 motifs, to characterize spatial signaling programs associated with mGAMs. Among the 20 discovered motifs (Supplementary Figures 15, 16, 17), two involved multicellular coordination between macrophages and tumor subtypes. In each motif, the communication was structured in a hub-and-spoke fashion with macrophages at the center. One of the two motifs had mGAMs at the center coordinating interactions with other mGAMs, multiple malignant tumor states (MES-like, AC-like, OPC-like, NPC-like), and vascular cells, with MES-like tumor cells as the dominant interaction partner (Figure 5c). In the second macrophage-centric motif, non-mGAM macrophages were instead at the center interacting with other non-mGAM macrophages and non-MES tumor subtypes (AC-like, OPC-like, NPC-like) (Figure 5f).

The two motifs contained notable similarities. In both, non-MES tumor cells were predominantly expressing ligands that were matched to receptors on the central macrophage subpopulation. The LRI families involved in signaling between the non-MES and macrophage were also consistent between the two, with near-equal contributions to total motif activity (Supplementary Figure 20a,b). Predominantly, non-MES tumor cells engaged in cell adhesion interactions, notably THY1 and JAM3 engaging integrin receptors ITGAM_ITGB2 and ITGAX_ITGB2, suggesting a role for early tumor cells in engaging macrophage through proximal junctions to localize them in the tumor microenvironment. Both motifs also show strong enrichment for tumor cells polarizing macrophage towards an M2-like state through the canonical CSF1-CSF1R axis.

Differences between the two motifs centered mostly on the additional MES-like interactions in the mGAM motif. MES-like-macrophage signaling was approximately 2.6-fold more concentrated in the mGAM motif than in the non-mGAM motif in both directions (Supplementary Figure 20c,d). The MES-like LRIs were enriched for many of the same interactions as the non-MES tumor cells (e.g. THY1, JAM3, CSF1, PDGFB), but at a lower relative level of enrichment (Fig. 5d). Several LRIs were enriched in the mGAM-MES-like interactions, in both directions, that were not seen highly enriched in any of the other tumor subtypes.

In particular, we observed enrichment of GRN-SORT1 in the mGAM→MES-like direction (Fig. 5d left). This enrichment is especially provocative as both GRN and SORT1 have substantial recent findings in GBM but have not been previously implicated together nor associated with specific macrophage subtypes. In multiple tumor types, macrophage-secreted granulin has been shown to remodel the tumor microenvironment and promote metastasis ^31,32^. In GBM mouse models, macrophage PGRN has been connected to tumor cell proliferation and mesenchymal protein expression ^33^, suggesting a potential causal role in transforming tumor cells to aggressive MES-like states. Parallel work in GBM cell lines has shown the Sortilin receptor is associated with invasion and mesenchymal transition ^34^. In human GBM data, a recent large-scale spatial multi-omic glioma atlas independently identified GRN as correlated with specific myeloid subpopulations in the glioma microenvironment^35^, providing orthogonal support for GRN as a biologically relevant macrophage-associated signal in human glioma.

In the MES-like→mGAM direction, we noted enrichment of ANXA1-FPR1 and TGF*β* signaling (Fig. 5d right). Expression of ANXA1 is a hallmark of mGAMs, yet it is also highly expressed in MES-like tumor cells.^26^ The connection between tumor-derived ANXA1 signaling and the mGAM subtype has not been explored. Recent work in high grade glioma has shown that ANXA1-FPR1 signaling can induce an M2-like state in microglia and macrophage. ^36^ Macrophage in high-ANXA1 tumors exhibited a more immunosuppressive phenotype and correlated with worse outcomes for patients, consistent with mGAM-related results.

We reasoned that the presence of the mGAM-centric motif may have clinical implications as it represents a potential shift towards a more aggressive glioma state. We first confirmed that the mGAM motif was enriched in high grade samples (Fig. 5e). Next, we used the ALARMIST downstream impacts for mGAMs present in this motif to assess potential directional changes of macrophages being transformed to mGAMs as a member of this motif. We constructed an mGAM signature using differentially expressed genes in these motif-active mGAMs. We selected the top 20 genes by log-fold change, all of which were significant at a false discovery rate of 0.001 (Fig. 5h). We used the gene set to construct a prognostic score for low grade glioma patients, the population at risk of transforming into an aggressive MES-like state which would then progress to GBM. Stratifying into high and low groups yielded a significant difference in clinical outcomes for LGG patients in TCGA (fig 5i).

Overall, the ALARMIST results provided biologically rational connections between existing observations of individual ligands and receptors in GBM and the recently-described mGAM cell type. Further, ALARMIST implicated specific ligands and receptors as potential drivers of MES-like transformation. The ALARMIST downstream impacts also validated the clinical relevance of the findings by discovering a gene score connected to worse outcomes in LGG patients.

## Discussion

To study recurrent spatial LRI patterns, we developed ALARMIST, a Bayesian Poisson matrix factorization framework for discovering coordinated cell–cell communication motifs from spatial transcriptomics data. Existing methods typically test individual ligand–receptor pairs independently and summarize spatial information at the cell type level, discarding the co-occurrence structure among signaling events within local tissue neighborhoods. ALARMIST jointly models all ligand–receptor interactions within local spatial units (patches), decomposing a matrix of LRI counts into latent communication motifs. Each motif captures a recurring, interpretable program of co-occurring signaling events enriched in specific microenvironments. ALARMIST inference leverages the sparsity of LRIs in each motif, leading to a computationally efficient method that can factorize large spatial LRI matrices in minutes on a laptop. We demonstrated the robustness of ALARMIST motif discovery and downstream impact estimation across both the 10x Xenium 5K and CosMx 6K platforms. We also demonstrated the ability of ALARMIST motifs to uncover novel biological insights and hypotheses in case studies of lung cancer progression and a malignant macrophage subtype in glioblastoma.

Like all transcriptomics-based cell-cell communication methods, ALARMIST assumes that mRNA levels correlate to protein abundance and localization. While evidence exists for moderate levels of correlation,^37^ this assumption can be violated by post-transcriptional regulation, protein trafficking, secretion dynamics, receptor internalization, and protein modifications such as phosphorylation, glycosylation, and proteolytic cleavage. ^38^ As in our LGG results, ALARMIST is best used as a hypothesis-generating analysis which then must be validated through direct measurement of protein expression.

The choice of patch size also influences results. A patch that is too large may mix distinct microenvironments; one that is too small may contain too few cells for reliable estimation. We default to 50*µ*m but recommend the user adjusts this if their tissue is sparse or a particular set of ligand-receptor interactions are known that warrant a different average scale of interaction. Relatedly, our binary classification of interactions into juxtacrine and paracrine is a simplification: secreted ligands diffuse with distance-dependent concentration gradients, whereas ECM components are deposited locally as insoluble scaffolds, yet both are modeled with the same spatial parameters. A natural extension would be to assign distinct spatial kernels to each interaction category. We also excluded autocrine signaling for juxtacrine interactions, as canonical pathways such as Notch-Delta require trans-interaction between distinct cells, while retaining it for ECM-receptor interactions where cells sense their own secreted matrix. These modeling choices provide reasonable defaults but could be refined as spatial multi-omics data and ligand diffusion estimates become more available.

Several extensions of the ALARMIST framework present promising directions. A hierarchical formulation could distinguish communication programs that are conserved across patients from those specific to individuals or tissue regions, enabling principled analysis of multi-patient cohort designs. The motif embeddings learned by ALARMIST also offer a basis for further cell subtyping: cells can be characterized not only by their transcriptional identity but by the communication programs they participate in, potentially revealing functional subtypes invisible to expression-based clustering alone. Refining the spatial modeling of different signaling modalities is another natural direction, including better profiling of paracrine, juxtacrine, and ECM-remodeling signals with category-specific distance models. As spatial profiling technologies continue to advance to provide more transcripts per cell, broader coverage of the transcriptome, and same-cell multiomics, we anticipate methods like ALARMIST will be increasingly needed to distill complex, high-dimensional read outs into compact and biologically interpretable signals.

## Methods

### Spatial transcriptomics datasets

Spatial transcriptomics datasets were profiled using the Xenium 5K platform (10x Genomics). We analyzed an in-house glioblastoma tissue microarray (TMA) dataset and a published lung adenocarcinoma progression cohort. The glioblastoma TMA comprised 13 punch cores from glioblastoma specimens (6 low-grade and 7 high-grade regions; *n* = 13 cores). The dataset contained 100,197 segmented cells profiled for 5,119 genes. Four patients contributed matched low-grade and high-grade cores (patients 9736, 23184, 19882 and 14007), enabling paired comparisons across tumor grades; patient 14007 additionally contributed replicate cores per grade. We selected four Xenium 5K samples from a published study of lung precursor lesion progression ^39^, consisting of matched adenocarcinoma in situ (AIS) and invasive lung adenocarcinoma (LUAD) from two patients (p17 and p21; *n* = 4 samples). Raw data were obtained from the study authors with permission for reanalysis.

### Data preprocessing and cell type annotation

For the GBM TMA dataset, segmented cells with fewer than 20 total detected transcripts were excluded. Cell types were annotated using a CellTypist model trained on the core GBMap reference atlas with annotation level 2. Within the myeloid compartment, we computed a *FOSL2* gene-module score using Scanpy; cells with module scores > 0.25 were labeled as mGAM, and the remaining myeloid cells were labeled as non-mGAM. Within the tumor compartment, cells were subtyped according to the cellular state framework of Neftel et al. ^30^. We obtained gene signatures for four GBM cellular states: mesenchymal-like (MES), astrocyte-like (AC), oligodendrocyte progenitor-like (OPC), and neural progenitor-like (NPC). Module scores were computed for each signature and converted to z-scores. Cells initially classified as differentiated-like were assigned to MES-like or AC-like based on which z-score was higher; cells classified as stem-like were similarly assigned to OPC-like or NPC-like.

Data from Ren et al. ^19^ were obtained from the SPATCH portal (https://spatch.pku-genomics.org/) and used for two analyses. For cross-platform concordance (Fig. 3), we used matched Xenium Prime 5K and CosMx 6K datasets, comprising consecutive tissue sections from one COAD, one OV, and one HCC specimen profiled on each platform (six sections total). For semi-synthetic benchmarking (Fig. 2), reference single-cell expression profiles were drawn from the matched COAD scRNA-seq dataset.

### Spatial patch construction

To quantify local cell-cell communication, each section was discretized into a regular grid of non-overlapping square patches with edge length *s* (in micrometers). In consideration of molecule diffusion and transcript bleeding in imaging-based platforms, we used *s* = 50 *µ*m unless otherwise specified. Cells were assigned to patches by their centroid coordinates; empty patches were excluded. For multi-sample analyses, LRI features were computed over the intersection of genes and the union of cell types shared across samples to ensure a common column space. Similarly, to construct spatial neighborhoods for each cell, neighborhoods were defined as a square patch of edge length *s* centered on that cell.

### Ligand-receptor interaction database

Ligand-receptor interactions (LRIs) were drawn from CellChatDB v2.0 or CellPhoneDB v5.0, filtered to pairs whose ligand and receptor genes were present in the dataset gene panel. Based on database annotations on interaction category (secreted signaling, ECM–receptor, or cell–cell contact), interactions annotated as cell–cell contact were assigned juxtacrine mode only. All remaining (non-contact) interactions were assigned paracrine mode; when the sender and receiver cell types are the same (*A* = *B*), an additional autocrine feature was created as a separate column in the LRI matrix, capturing co-expression of ligand and receptor within individual cells.

### LRI counting

To count LRI events, gene expression was binarized: a gene was considered expressed if its raw transcript count exceeded zero. This binary representation is appropriate for imaging-based spatial transcriptomics platforms (e.g., Xenium), where transcript detection is sparse and count magnitudes are less informative than in sequencing-based methods, making LRI counts less sensitive to technical variation in detection efficiency. Each LRI feature is indexed by a sender cell type *A*, receiver cell type *B*, ligand complex *L*, receptor complex *R*, and signaling mode *m* ∈ {autocrine, paracrine, juxtacrine}:

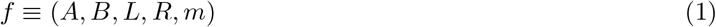

Within a given patch *i*, we assume that any ligand-expressing cell can signal to any receptor-expressing cell regardless of signaling mode. Even for contact-dependent (juxtacrine) interactions, we do not further restrict to immediately adjacent cells, as transcript bleeding in imaging-based platforms blurs the spatial precision of expression assignments at the sub-patch scale. For a given patch *i* and feature *f*, let *n*_*L*_ denote the number of type-*A* cells expressing the ligand and *n*_*R*_ the number of type-*B* cells expressing the receptor. For single-gene ligands or receptors, this is simply the count of expressing cells. Many ligands and receptors, however, are multi-subunit complexes requiring co-expression of all constituent genes for functional signaling. Let *L* = {*l*_1_, …, *l*_*m*_} and *R* = {*r*_1_, …, *r*_*n*_} denote the subunit genes of the ligand and receptor. We approximate the number of complex-expressing cells as the minimum count across subunits:

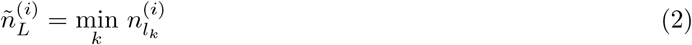

and analogously for the receptor. The interaction count is then computed according to the signaling mode.

#### Autocrine

A cell expresses both ligand and receptor, enabling self-stimulation. This mode applies only to non-juxtacrine LRIs when *A* = *B*. We apply strict AND logic at the cell level: *C*_auto_ equals the number of type-*A* cells co-expressing all ligand and receptor genes.

#### Paracrine

Distinct cells serve as sender and receiver. The interaction count is

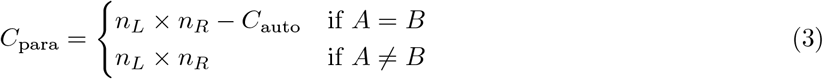

where the subtraction when *A* = *B* avoids double-counting cells that express both ligand and receptor.

#### Juxtacrine

Contact-dependent LRIs (e.g., Notch–Delta) do not admit autocrine signaling. The interaction count is

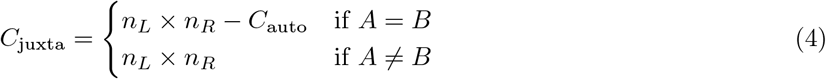

The product *n*_*L*_ × *n*_*R*_ represents the number of possible sender-receiver cell pairs, treating each pair as a candidate signaling event. This counting approach yields non-negative integer values that naturally suit Bayesian Poisson tensor factorization. After constructing the patch–LRI count matrix, features with zero counts across all patches were removed.

### Bayesian Poisson tensor factorization

The resulting counts across all patches form a matrix 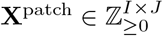, where *I* is the number of patches and *J* the number of LRI features. To discover recurring patterns of LRI co-activity across tissue regions, we decompose **X**^patch^ using Bayesian Poisson tensor factorization (BPTF). BPTF approximates **X**^patch^ as the product of two non-negative factor matrices:

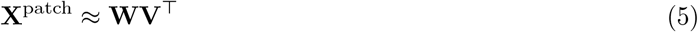

where 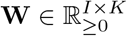 contains patch loadings and 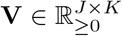 contains LRI factor loadings, with *K* denoting the number of latent motifs. Each entry is modeled as

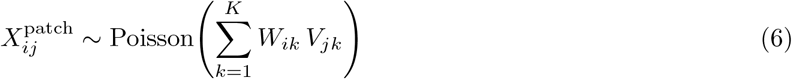

with gamma priors on both factor matrices:

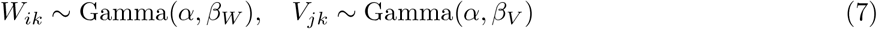

where *α* is shared across both modes and *β*_*W*_, *β*_*V*_ are mode-specific rate parameters. Inference follows Schein et al. ^18^. We use their implementation with default hyperparameters and set the number of motifs *K* as a user-specified parameter (see dataset-specific parameters below).

### Factor normalization and interpretation

Raw LRI factors from BPTF reflect both motif-specific patterns and the overall prevalence of each LR pair in the dataset. To prevent ubiquitously abundant LRIs from dominating motif characterization, we normalize by global LR pair prevalence. For each LRI feature *j* associated with ligand–receptor pair (*L, R*), we define the global prevalence *µ*_(*L,R*)_ as the sum of per-feature mean counts over all LRI features sharing the same ligand and receptor genes (aggregating across cell type combinations and signaling modes). The prevalence-normalized factor is then:

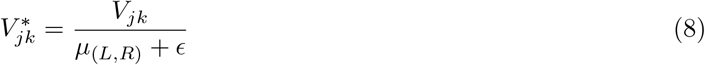

where *ϵ* = 1 serves as a smoothing constant. High values of 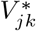 indicate LRIs specifically enriched in motif *k* relative to their background. The model is scale-invariant: *W*_*ik*_ → *W*_*ik*_*/c*_*k*_, *V*_*jk*_ → *V*_*jk*_ · *c*_*k*_ leaves the reconstruction unchanged. We set *c*_*k*_ = max_*i*_ *W*_*ik*_, yielding

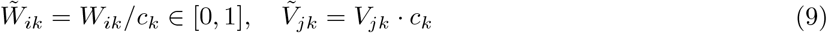

where 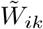 gives the relative activation of motif *k* across patches, and 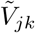 absorbs the motif-specific magnitude for cross-motif comparison. For downstream motif characterization, we report the combined score 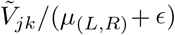, which reflects both the motif-specific enrichment and the absolute magnitude of the factor.

### Single-cell projection

The patch loading matrix **W** summarizes motif activity at the patch level, averaging over all cells within each neighborhood. To resolve motif activation at single-cell resolution, ALARMIST projects the patch-level factorization to individual cells. For each cell, we construct a spatial neighborhood with the same edge length *s* as the patches used for motif discovery and compute LRI features using the identical counting procedure, yielding a cell-LRI matrix 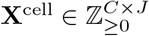 with the same column structure as the patch-level matrix. We then fix the learned LRI factors **V** from the patch-level model and infer cell loadings 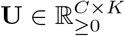 by updating only the cell loading parameters while holding **V** clamped:

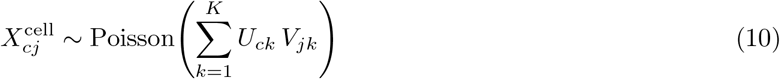

The resulting cell loadings *U*_*ck*_ measure how strongly the local environment of cell *c* expresses motif *k*. Cells are processed in chunks (default 50,000 cells) to maintain bounded memory usage.

### GMM-based motif state calling

The continuous cell loadings provide graded motif activation, but downstream analyses often require discrete assignments. ALARMIST classifies each cell as motif-positive or motif-negative using a two-component Gaussian mixture model (GMM) on log-transformed cell loadings. For each motif *k*, the log transformation separates the typically right-skewed loading distribution into two modes. The component with the higher mean is designated as the positive state, and each cell is classified by maximum a posteriori assignment. The posterior probability of the positive state is retained as a continuous confidence score. For multi-sample analyses, the GMM is fit on pooled cell loadings across all samples to ensure consistent thresholds.

### Motif–gene association via Poisson GLM

To investigate whether motif activity is associated with gene expression changes beyond the defining LRI genes, we fit a Poisson generalized linear model (GLM) without regularization for each non-LR gene *g*, motif *k*, and cell type *t*:

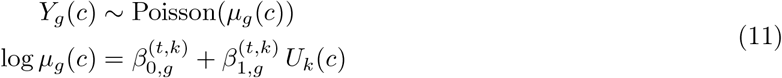

where *Y*_*g*_(*c*) is the raw transcript count of gene *g* in cell *c*, and *U*_*k*_(*c*) is the z-scored log motif loading of motif *k*, standardized within cell type *t*. Cells with zero motif loadings were excluded from model fitting. A positive 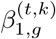 indicates gene upregulation with increasing motif activity; a negative value indicates downregulation. Significance is assessed by Wald test with Benjamini–Hochberg FDR correction (*α* = 0.05). All ligand and receptor genes contributing to the LRI features are excluded from testing to avoid circularity. To accelerate computation, ALARMIST optionally applies a Spearman rank correlation pre-filter (uncorrected *p* < 0.001) before fitting the full Poisson GLM, with FDR correction applied to the filtered gene set. Imaging-based spatial transcriptomics platforms are subject to transcript bleeding, where transcripts from neighboring cells are misassigned to the focal cell. To mitigate this artifact, we identify marker genes for each cell type by Mann–Whitney U test on normalized log-transformed expression (Benjamini–Hochberg adjusted *p* < 10^−5^, log_2_ fold change > 1) and exclude markers of other cell types when reporting results for cell type *t*.

### Cross-platform concordance of LRI motifs and spatial niches

To assess cross-platform robustness, we applied ALARMIST to matched Xenium 5K and CosMx 6K datasets of COAD, OV, and HCC from Ren et al.Ren et al.^19^, where consecutive tissue sections were profiled on each platform. All samples were processed with identical parameters (50 *µ*m patch radius, *K* = 20).

#### Spatial coordinate alignment

Because serial sections do not share a coordinate system, we aligned CosMx cell coordinates to the Xenium coordinate frame. Corresponding anatomical landmarks (≥3 pairs per disease) were selected from tissue density heatmaps, and a similarity transform (rotation, uniform scaling, translation) was estimated by least-squares optimization. The transform was applied to all CosMx cell coordinates.

#### LRI factor comparison

For each cancer type, we computed pairwise cosine similarities between Xenium and CosMx motif LRI factor vectors (columns of **V**), restricted to cell-type–LRI combinations present on both platforms. A motif was classified as well-aligned if its maximum cosine similarity with any motif on the opposing platform exceeded 0.4, and poorly aligned otherwise. For each Xenium motif, the CosMx motif with the highest cosine similarity (≥ 0.4) was designated as its matched counterpart. To diagnose sources of discordance, we extracted the top 50 LRIs per motif (ranked by LRI factor value) and computed the overlap rate with the opposing platform’s pooled effective LRI set at three levels: individual LR genes, LR pairs, and cell-type–LRI combinations.

#### Spatial concordance of matched motifs (Lee’s *L*)

To quantify whether matched motif pairs co-localize across platforms, we computed Lee’s *L* statistic on the overlapping field of view. The intersection of the two platforms’ bounding boxes was divided into spatial bins of side length 100 *µ*m. For each matched motif pair, we computed the proportion of motif-positive cells per bin on each platform, yielding vectors **x** and **y** over *n* valid bins. Define the spatial lags 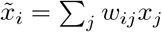 and 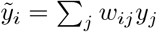. Lee’s *L* is given by:

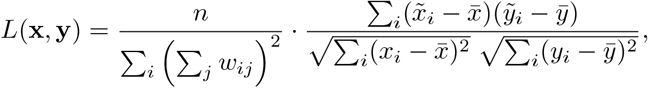

where **W** is a row-normalized queen contiguity spatial weights matrix on the bin grid (*w*_*ij*_ > 0 if bins *i* and *j* share an edge or corner). Bins with fewer than one cell from either platform were excluded, and a minimum of 20 valid bins was required.

#### Niche clustering and concordance

For each sample, cell-level motif loadings were restricted to matched motifs and normalized per motif by the maximum loading. *K*-means clustering was applied with *K* determined by elbow analysis on the Xenium sample; the same *K* was used for CosMx. To compare niches across platforms, the overlapping field of view was divided into 500 *µ*m grid squares (excluding squares with < 10 cells from either platform). For each grid square *g*, we computed the niche proportion vector:

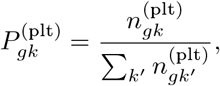

where 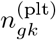 is the number of cells assigned to niche *k* in grid square *g* on platform plt. Each CosMx niche *j* was matched to the Xenium niche 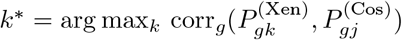, and the corresponding Pearson *r* was reported as the per-niche concordance score.

#### Downstream impact concordance

For each cancer type, per-gene spatial expression correlation was computed by normalizing expression matrices (total-count, log1p), computing pseudo-bulk means per cell type, and calculating Pearson *r* per gene across shared cell types. Downstream impact was assessed using GLM log-fold changes from ALARMIST’s impact analysis. For each matched motif pair and shared cell type, genes significant on either platform (*q*< 0.05) were retained. Genes were divided into 10 quantile bins by spatial expression correlation. Within each bin, Pearson *r* of CosMx versus Xenium log-fold changes was computed with 95% confidence intervals from 1,000 bootstrap resamples. A linear regression was fit to the binned values.

#### Spatial autocorrelation of gene expression

Moran’s *I* was computed per gene per sample using squidpy’s spatial_autocorr with a KNN spatial graph (*n*_neighbors_ = 20) on normalized log-transformed expression.

### Dataset-specific parameters

For the GBM TMA dataset, ALARMIST was run with patch size *s* = 50 *µ*m using CellChatDB v2.0. BPTF was fit with *K* = 20 motifs. To compare motif activity between low-grade and high-grade glioblastoma regions, we computed the fraction of motif-positive cells (as determined by GMM state calling) of each TMA core. Differences in motif-positive fractions between grades were assessed using the Mann–Whitney U test across cores (*n* = 7 per grade). For the LUAD dataset, ALARMIST was run with patch size *s* = 80 *µ*m using CellChatDB v2.0. BPTF was fit with *K* = 25 motifs. To characterize transcriptional programs associated with disease progression, we defined three mutually exclusive cell groups based on motif state assignments: cells positive for healthy vasculature motif only, cells positive for tumor vasculature motif only, and cells positive for both motifs. Gene module scores for Hallmark pathway gene sets (MSigDB) were computed using `sc.tl.score_genes` on normalized log-transformed expression and compared across the three groups within each cell type using Mann–Whitney U tests. Pairwise differential expression analysis between motif groups was performed within each cell type using the Mann–Whitney U test on per-sample normalized and log-transformed expression, with Benjamini–Hochberg FDR correction. Gene set enrichment analysis (GSEA) was performed on the log_2_ fold change-ranked gene lists using gseapy prerank against the MSigDB Hallmark 2020 collection (1,000 permutations; FDR < 0.25).

### Semi-synthetic design

The goal of our semi-synthetic design is two-fold: to evaluate ALARMIST’s ability to recover motifs and detect significantly upregulated and downregulated genes in a setting where the ground-truth is known. We repeat the entire procedure, generating the simulated data and fitting each method to it, ten times.

#### Spatial structure

For each motif *m*, we sample *Z*_*m*_, a 2D Gaussian process over the spatial domain with kernel

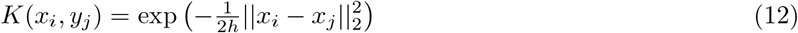

where *h* = 0.09. We then sample *q* ∼ Uniform(0.6, 0.9) and define the active region as the set {(*x, y*) : *Z*_*m*_(*x, y*) > Quantile(*Z*(*x, y*), *q*)}, which is constrained to cover between 10-40% of the domain. The corresponding spatial regions are shown in Fig. 2. We then sample another Gaussian process to sample cells in the space. In particular, for each cell type, we sample *Z*_*ct*_ ∼ GP, *h* = 0.01 according to an inhomogeneous Poisson point process proportional to the sum between the exponentiated surface induced by the Gaussian process, and indicators of motif activity. In particular,

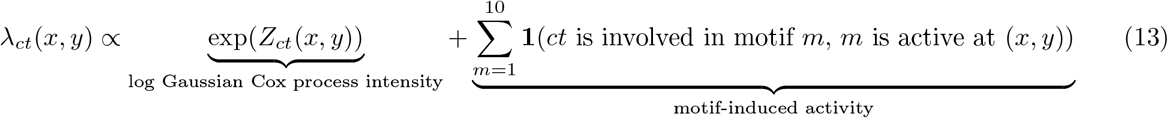

where *Z*_*ct*_(*x, y*) is the GP surface at location (*x, y*). We scale this term so that each patch, on average, contains 25 cells. We then sample gene expression vectors for each cell location with replacement from the vectors corresponding to that cell type in the reference data.

#### Modifying gene expression, conditional on the regions

We augment the cell by gene matrix *Y* accordingly. For each cell *c* and gene *g*: if the cell is active in a program and its cell type *ct* is in the cell type interaction network, and the gene is a corresponding active gene: resample the gene expression, sampling *q* ∼ Beta(9, 1) and assign *Y*_*g*_(*c*) = Quantile_*g,ct*_(*q*) the *q*th quantile of that cell type’s expression level in the reference data. If it is a downregulated gene, we sample *q* ∼ Beta(1, 9). If the cell-gene combination is present in the motif, but the cell lies in an inactive region, then we also sample *q* ∼ Beta(1, 9). If the cell-gene combination does not appear in any of the motifs, we instead binomial thin the value, sampling 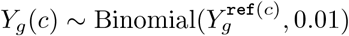. This final step reduces the existing structure in the reference COAD data, which for the purposes of this experiment, we are uninterested in.

We vary the size of the spatial domain to yield {100, 400, 1000, 5000, 10, 000} patches of size 50 × 50 and scale the inhomogeneous Poisson process intensity so that on average, there are approximately 25 cells per patch.

#### Sender-receiver network recovery

We first evaluate ALARMIST’s ability to recover each motif’s cell type network (shown in Fig. 2b). We consider ALARMIST, Tensor-cell2cell, COMPOTES, and NICHES, using the tutorials in each method’s respective Github repository to run and analyze the benchmarks. We fit each method with 20 components, providing an allowance of 10 extra components to capture existing noise or redundancy in the data. ALARMIST, Tensor-cell2cell and COMPOTES yield (sender cell type, receiver cell-type, motif) tensors which may be reconstructed directly from the estimated parameters of each method. We matricize each into a 16^2^ × 20 matrix and match the estimated interaction networks to their closest ground truth ones using the Hungarian matching algorithm to maximize the mean cosine similarity across motifs. Since NICHES does not directly provide a cell type interaction network, but instead a cell-cell signaling matrix, and we apply K-means clustering (*K* = 20 clusters) to the columns of the cell-cell signaling matrix after standard pre-processing the columns (log-transform, normalization and standardization) and dimensionality reduction via PCA (choosing 30 principal components). We then aggregate by sender and receiver cell type to obtain the corresponding cell type interaction by program matrix for NICHES.

#### Impact

Alarmist provides cell loadings; we use these cell loadings to identify motif-associated gene expression changes using the Poisson GLM framework described above. We compare to two hard-clustering based methods, Leiden clustering at resolution levels (0.1, 0.5, 1.0), and a recently introduced spatial foundation model, Novae ^20^. After finetuning Novae to the synthetic data *Y*, we use Novae to obtain cell embeddings and ran K-means (K=10, equal to the ground-truth) on the cell embeddings to form clusters. We also apply Louvain clustering using Seurat’s `FINDCLUSTERS` function, with resolution parameter set to 0.5, on the global NICHES neighborhood-to-cell signaling matrix. This procedure yields clusters of receiving cells. For each method and cell type, we match clusters to the ground truth motifs using Jaccard similarity. For the hard clustering methods, we use a dummy variable approach to indicate cluster membership rather than the log cell embedding, as is standard for clustering methods.

Out of the significant genes, we select the ten non-LRI genes with the highest absolute log-fold change. Comparing these sets to the ground truth, also consisting of ten marker genes, we compute the F1 score, defined as 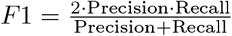. In this setting, since there are ten selected genes and ten ground-truth genes per motif, the F1 score simplifies to 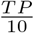.

#### Significant differences in F1

To determine significant differences in motif-specific F1 scores between ALARMIST and the baseline methods, we run Wilcoxon signed-rank tests ^40^. The Wilcoxon signed-rank test is a nonparametric statistical test for testing whether the median of the difference of two sets of matched samples deviates from zero.

#### Lee’s L

For each spatial plot in Fig. 2g and Fig. 2h, Lee’s L is a bivariate spatial correlation coefficient which measures the association between two sets of observations made at the same locations. For pairs of observations 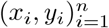, where *x*_*i*_ and *y*_*i*_ occur at the same location, define 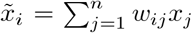 and 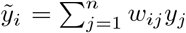. Then Lee’s L is given by

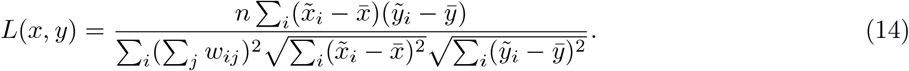

To construct *W*, we define 𝒩 (*i*) to be the set of the 50 nearest neighbors, as measured by Euclidean distance, to point *i*. Then

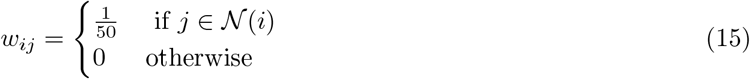

## Code availability

ALARMIST is available as an open-source Python package at https://github.com/tansey-lab/alarmist. All analyses and figure generation scripts used in this study are provided in the same repository under https://github.com/tansey-lab/alarmist/blob/main/tutorials/GBM.ipynb. ^1^

## Code availability

All ALARMIST code is publicly available on GitHub (https://github.com/tansey-lab/alarmist).

## Data availability

The CosMx 6K and Xenium 5K datasets of COAD, OV, and HCC were obtained from Ren et al.Ren et al.^19^. The Xenium 5K datasets of AIS and LUAD were obtained from Peng et al. ^22^. Accession numbers and download links are provided in the corresponding original publications. The Xenium 5K glioma dataset will be made publicly available upon publication.

## Acknowledgments

WT is supported by the NIH/NCI (R37 CA271186, U54 CA274492, P30 CA008748), Break Through Cancer, the Fund for Innovation in Cancer Informatics, the Cancer AI Alliance, the Tow Center for Developmental Oncology, and the Maurice Campbell Initiative at Memorial Sloan Kettering Cancer Center. JH is supported by the NSF under Grant No. 2140001.

## Author Information

### Author contribution

J.F. led the development and implementation of the model and the analysis of datasets. J.H. designed and performed benchmarking analyses. J.S. performed co-culture experiments. J.F.Q. assisted with code organization and Nextflow pipeline integration. Y.D. contributed to the analysis and interpretation of the AIS and LUAD datasets. A.S. and W.T. conceived the project and contributed to the development of the Bayesian Poisson tensor factorization framework. K.K.H.Y. provided clinical and biological expertise on glioblastoma and guided the formulation of biological questions. W.T. oversaw the project and guided the development of the model and interpretation of the results. Members of the Break Through Cancer Data Science TeamLab reviewed the manuscript and contributed to internal discussions during development of the method.

### Full list of Data Science Hub TeamLab

Cheng-Zhong Zhang, Ethan Cerami, Franziska Michor, Nathalie Agar, Rameen Beroukhim, Ashley Kiemen, Atul Deshpande, Dimitri Sidiropoulos, Genevieve Stein-O’Brien, Leslie Cope, Rachel Karchin, Roy Elias, Kadir Akdemir, Linghua Wang, Charlie Whittaker, Stuart Levine, Andrew McPherson, Benjamin Greenbaum, Niki Schultz, Sohrab Shah, Elana Fertig, Renad Al Ghazawi, Nil Aygun, Archana Balan, Gerard Baquer, Vincent Butty, Nick Ceglia, Jennifer Chen, Yu-Chen Cheng, Andrew Cherniack, Kate Cho, Kyung Serk (Kevin) Cho, Chase Christenson, Simona Cristea, Yibo Dai, Simona Dalin, Ino de Bruijn, Charlie Demurjian, Henry Dewhurst, Huiming Ding, Tom Dougherty, Emma Dyer, Yuval Elhanaty, Jiayi Fan, Sasha (Alexander) Favorov, Andre Forjaz, Samuel Freeman, Alessandro Grande, Christopher Graser, Benjamin Green, Eliyahu Havasov, Jason Hwee, Afrooz Jahedi, Jiahui Jiang, Taisha Joseph, Saurabh Joshi, Nancy Jung, Luciane Kagohara, Jennifer Karlow, Jiaying Lai, Kevin Meza Landeros, Joshua Lau, Nora Lawson, Jake Lee, Hannah Lees, Matt Leventhal, Alex Ling, Lester (Yunzhou) Liu, Yang Liu, Yunhe Liu, Dmitrijs Lvovs, Valentina Matos Romero, Marcus Mendes, Jose Meza Llamosas, Thomas Mitchell, Hideaki Mizuno, Jacob Myers, Matthew Myers, Nataly Naser Al Deen, Siri Palreddy, Sergiu Pasca, Jeffrey Quinn, Sabahat Rahman, Kimal Rajapakshe, Shriya Rangaswamy, Greg Raskind, Shahab Sarmashghi, Manuel Schürch, Alyza Skaist, Alexander Solovyov, Haruna Tomono, Michael Toomey, Chris Tosh, Mesut Unal, Yann Vanrobaeys, Miriam Vines, Jun Wang, Meredith Wetzel, Marc Williams, Juan Xie, Lingqun Ye, Long Yuan, Syed Zaidi, Matthew Zatzman, Haoran Zhang, Bo Zhao, Peter Zheng, Feiyang Huang, Irika Katiyar, Elana Sverdlik.

## Footnotes

1 Due to the computationally expensive nature of the semi-synthetic simulations, we separately provide the Python scripts for these experiments.

