## Supplementary Figures for "Decoding Multicellular Communication Motifs from Spatial Transcriptomics with ALARMIST"

### A Supplemental

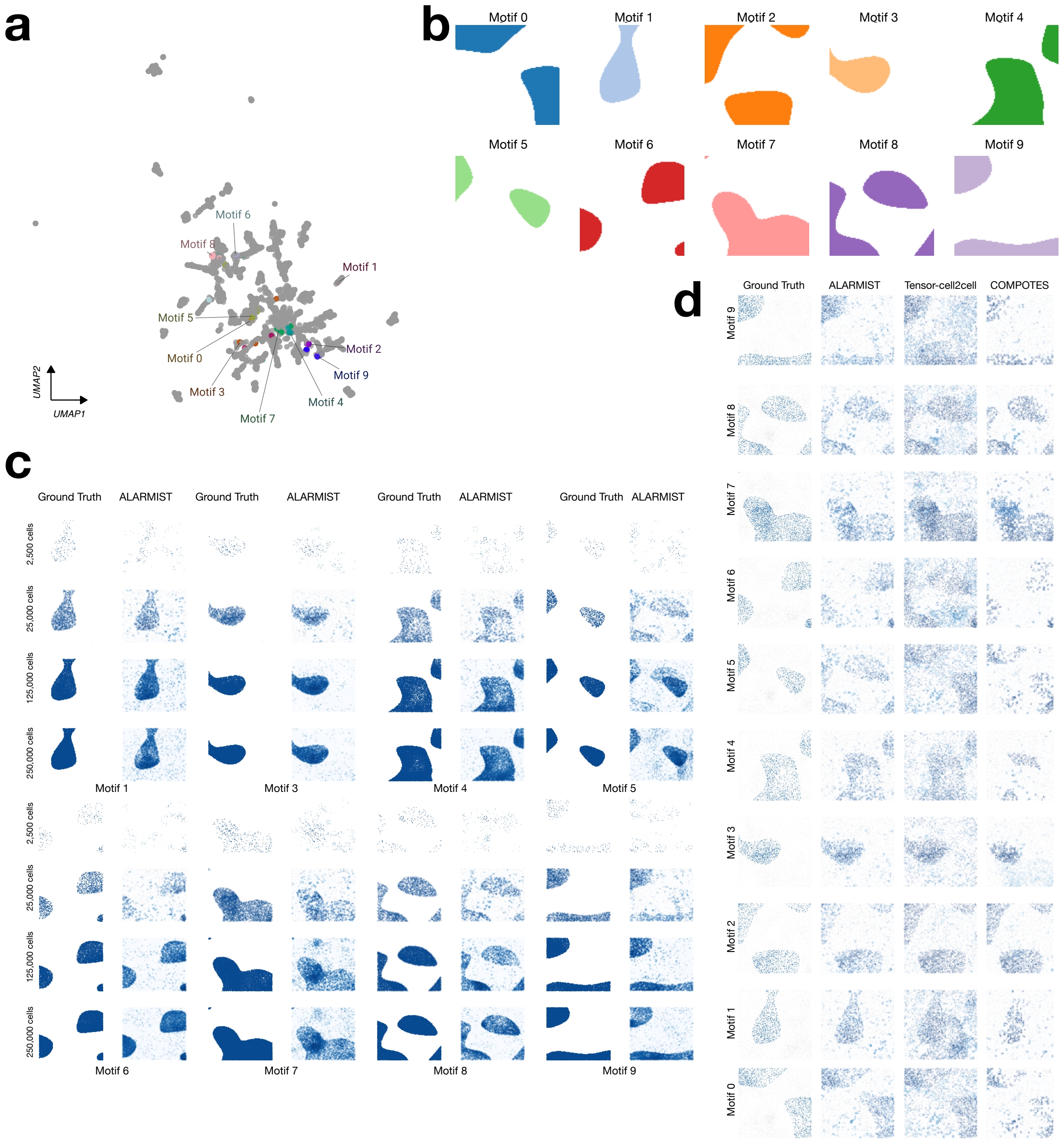


**Supplementary Figure 1: Additional benchmark results.** a. UMAP of all LRI factors. b. Each motif's active spatial region. c. Additional spatial motif recovery comparisons. d. Additional spatial comparisons of ALARMIST for different numbers of cells.

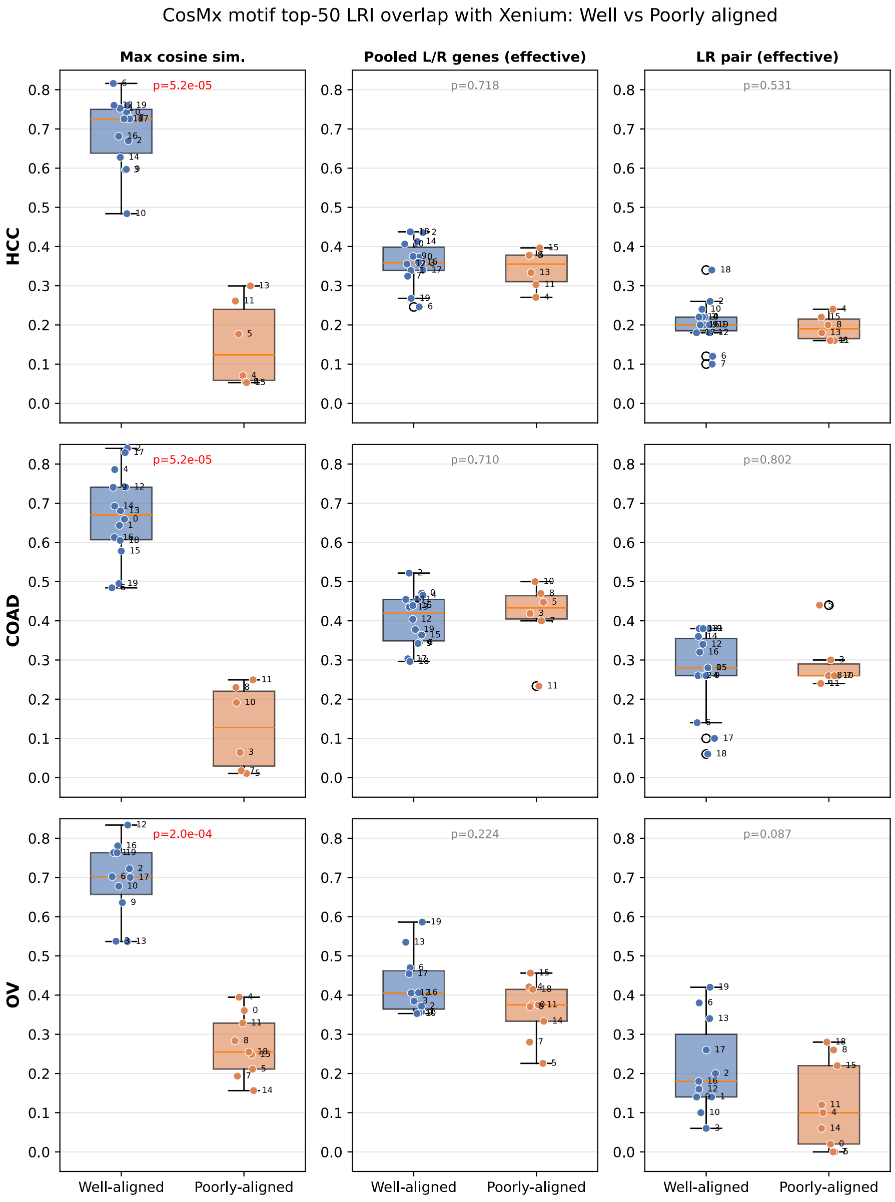


**Supplementary Figure 3: LRI overlap between CosMx motifs and the Xenium effective LRI pool, stratified by alignment status.** Each row corresponds to one cancer type (HCC, COAD, OV). Each CosMx motif is classified as well-aligned (blue) or poorly aligned (orange) based on whether its maximum cosine similarity with any Xenium motif exceeds 0.4. For each CosMx motif, the top 50 LRIs (ranked by LRI factor value) were compared to the pooled top-50 LRIs across all Xenium motifs. **Left column:** Maximum cosine similarity with Xenium motifs (confirmation of group assignment; significant by construction). **Middle column:** Overlap rate at the level of individual ligand and receptor genes. **Right column:** Overlap rate at the level of LR pairs. No significant difference was observed at the gene or LR-pair level ($p > 0.05$, two-sided Mann–Whitney $U$ test), indicating that both groups of CosMx motifs use similar genes and LR pairs. The significant difference emerges only at the cell-type–LRI level (Fig. **??**c), where cell-type identity is included. Each dot represents one CosMx motif, labeled by motif index.

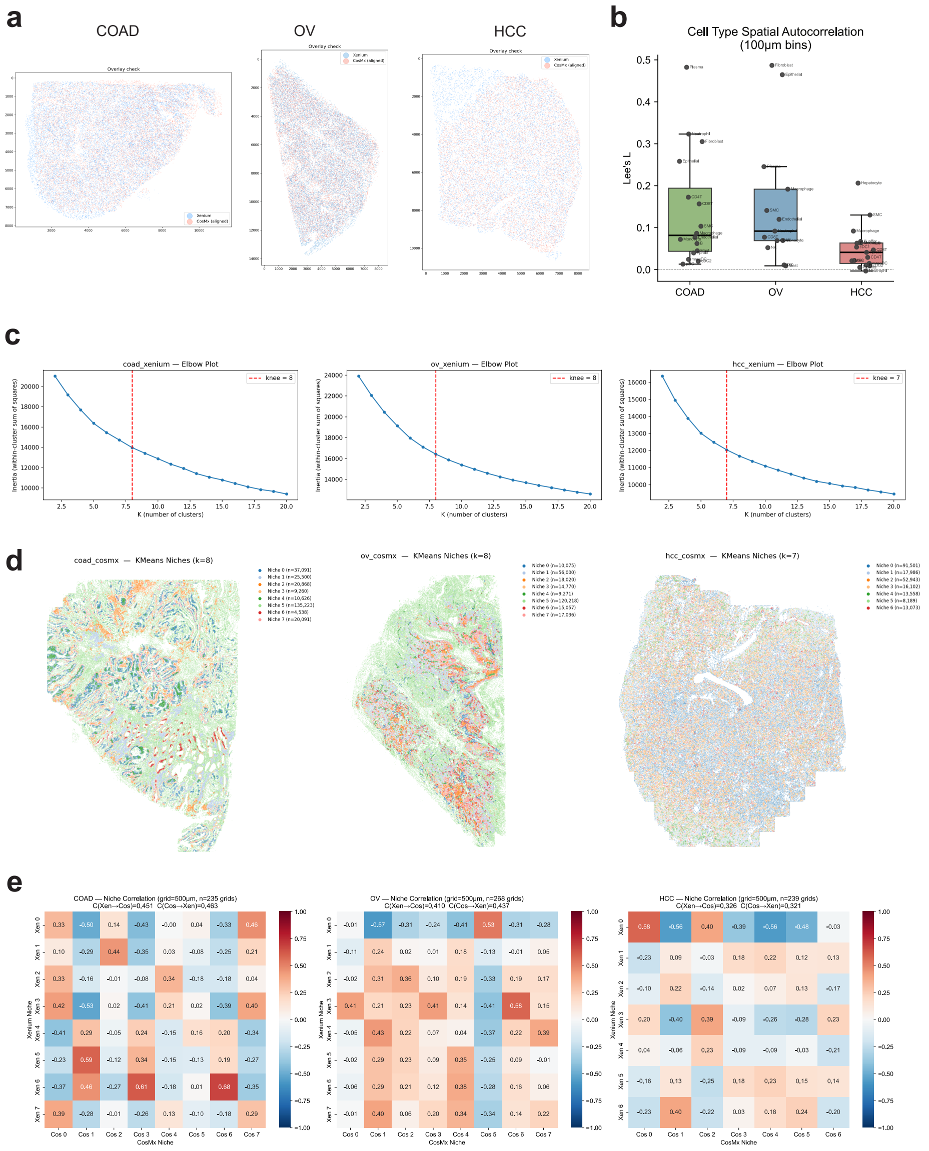


**Supplementary Figure 4: Cross-platform niche concordance: registration, clustering, and correlation details.** **a.** Spatial overlay of Xenium (blue) and CosMx (red) cell coordinates after similarity-transform alignment for COAD, OV, and HCC. Overlap confirms registration quality across all three cancer types. **b.** Lee's $L$ of cell-type proportions across platforms, computed per cell type on 100 $\mu$m bins with queen contiguity spatial weights (same setup as Fig. **??**d). Each dot represents one shared cell type, labeled by name. Cell-type Lee's $L$ values are generally comparable to or lower than matched-motif Lee's $L$ values (Fig. **??**d), indicating that ALARMIST motifs capture cross-platform spatial concordance beyond cell-type composition alone. **c.** Elbow plots for $K$-means niche clustering on Xenium samples. Inertia (within-cluster sum of squares) is shown as a function of $K$. Red dashed line indicates the selected $K$ (COAD: 8, OV: 8, HCC: 7), determined by the kneedle algorithm. The same $K$ was applied to the corresponding CosMx sample. **d.** Spatial maps of $K$-means niche clusters for CosMx samples, colored by cluster identity. Colors are not matched to Xenium niches; see Fig. **??**e for the matched-color version. **e.** Pearson correlation matrices of niche proportion vectors between Xenium (rows) and CosMx (columns) across 500 $\mu$m grid squares for each cancer type. Entry $R_{kj}$ is the Pearson correlation between the proportion of Xenium niche $k$ and CosMx niche $j$ across all valid grid squares. Summary concordance scores $C(\text{Xen} \rightarrow \text{Cos})$ and $C(\text{Cos} \rightarrow \text{Xen})$ (mean best-match correlation) are reported in each panel title.

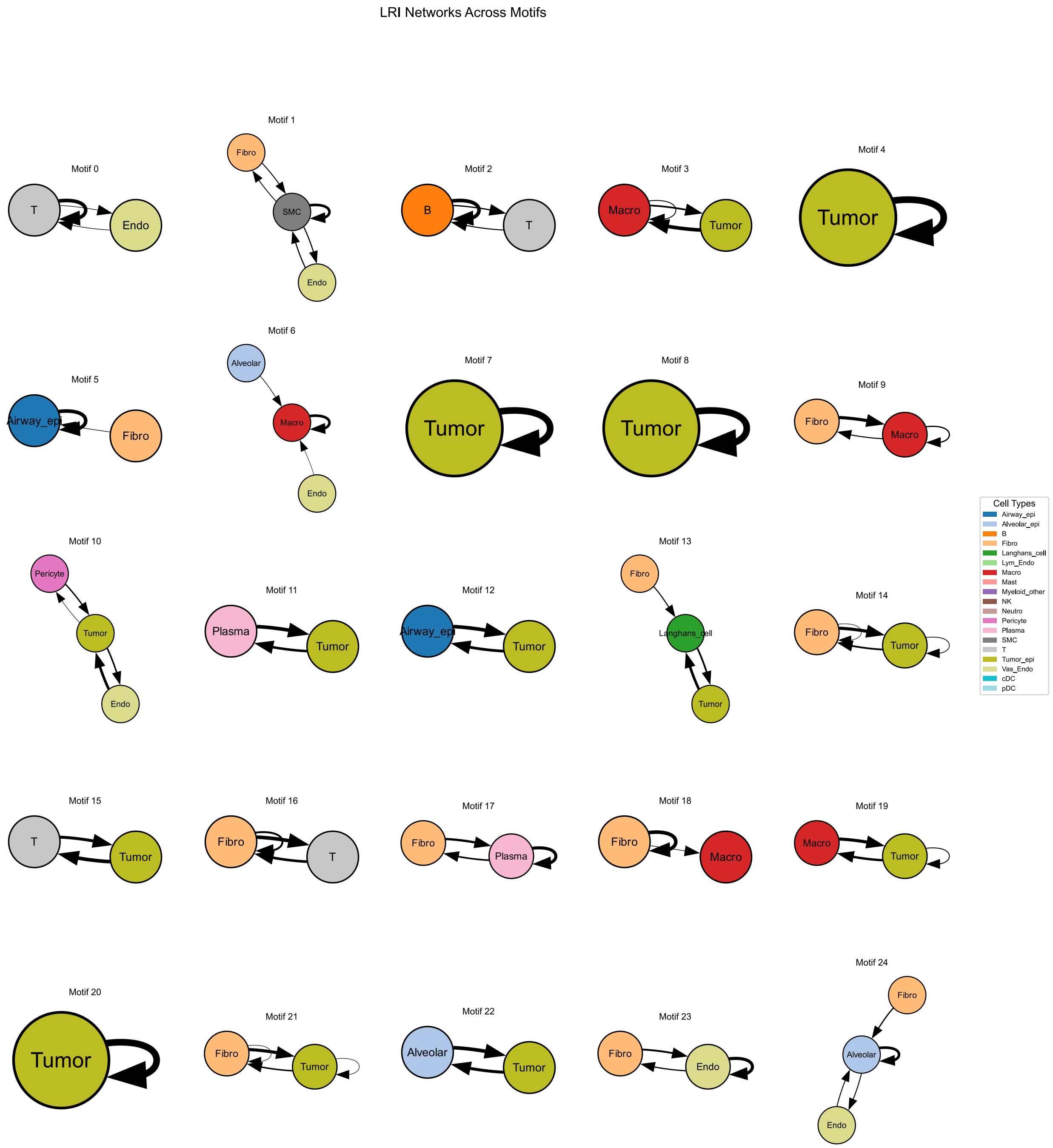


**Supplementary Figure 5: Cell-type interaction networks for all 25 LUAD motifs.** Each panel shows one motif as a network graph. For each motif, the top 500 LRIs (ranked by normalized factor value) were aggregated by sender–receiver cell-type pair. Cell types with aggregated factor values exceeding 800 are displayed as nodes, with edges representing LRI interactions between them. Edge thickness reflects the summed factor value.

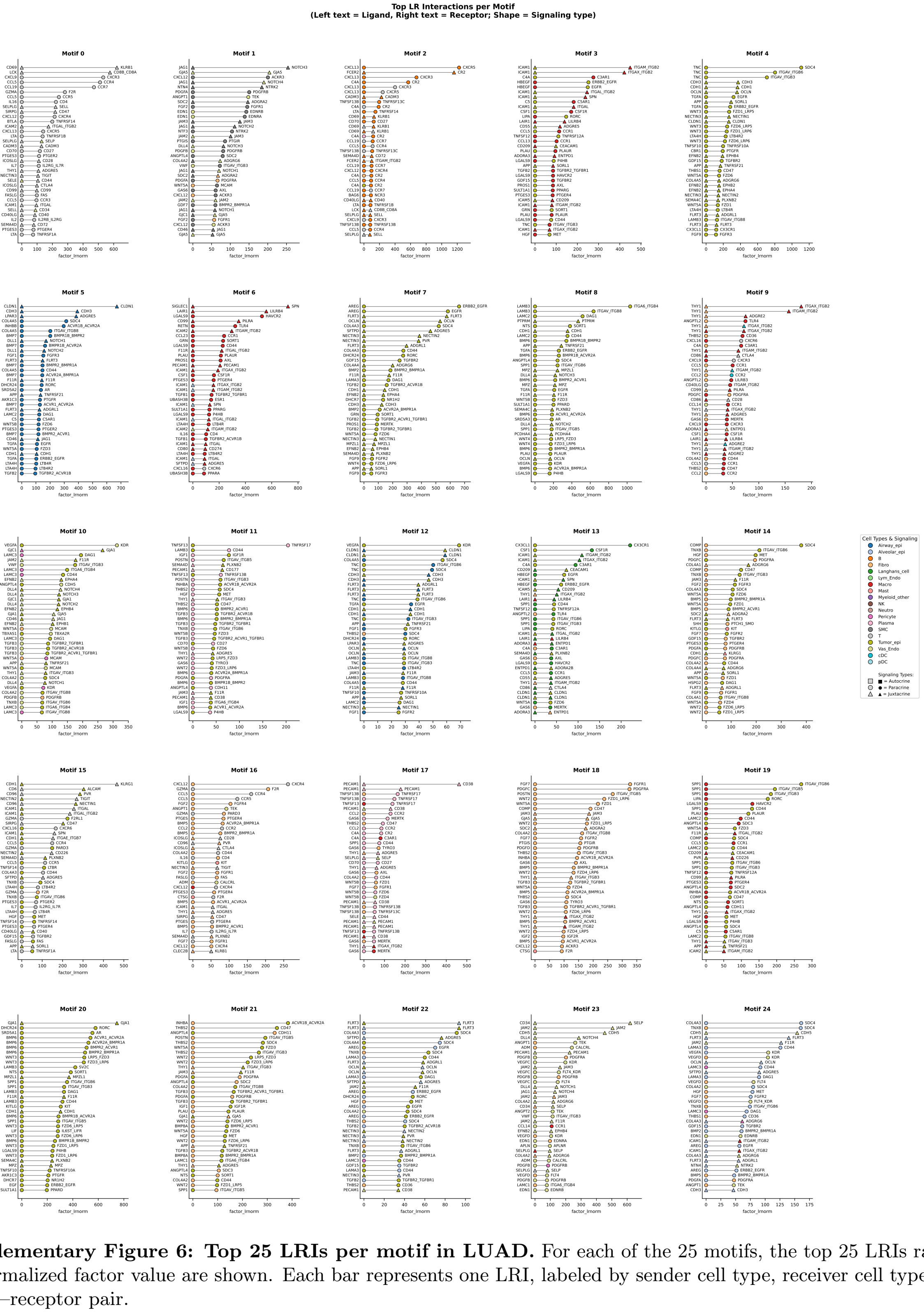


**Supplementary Figure 6: Top 25 LRIs per motif in LUAD.** For each of the 25 motifs, the top 25 LRIs ranked by normalized factor value are shown. Each bar represents one LRI, labeled by sender cell type, receiver cell type, and ligand–receptor pair.

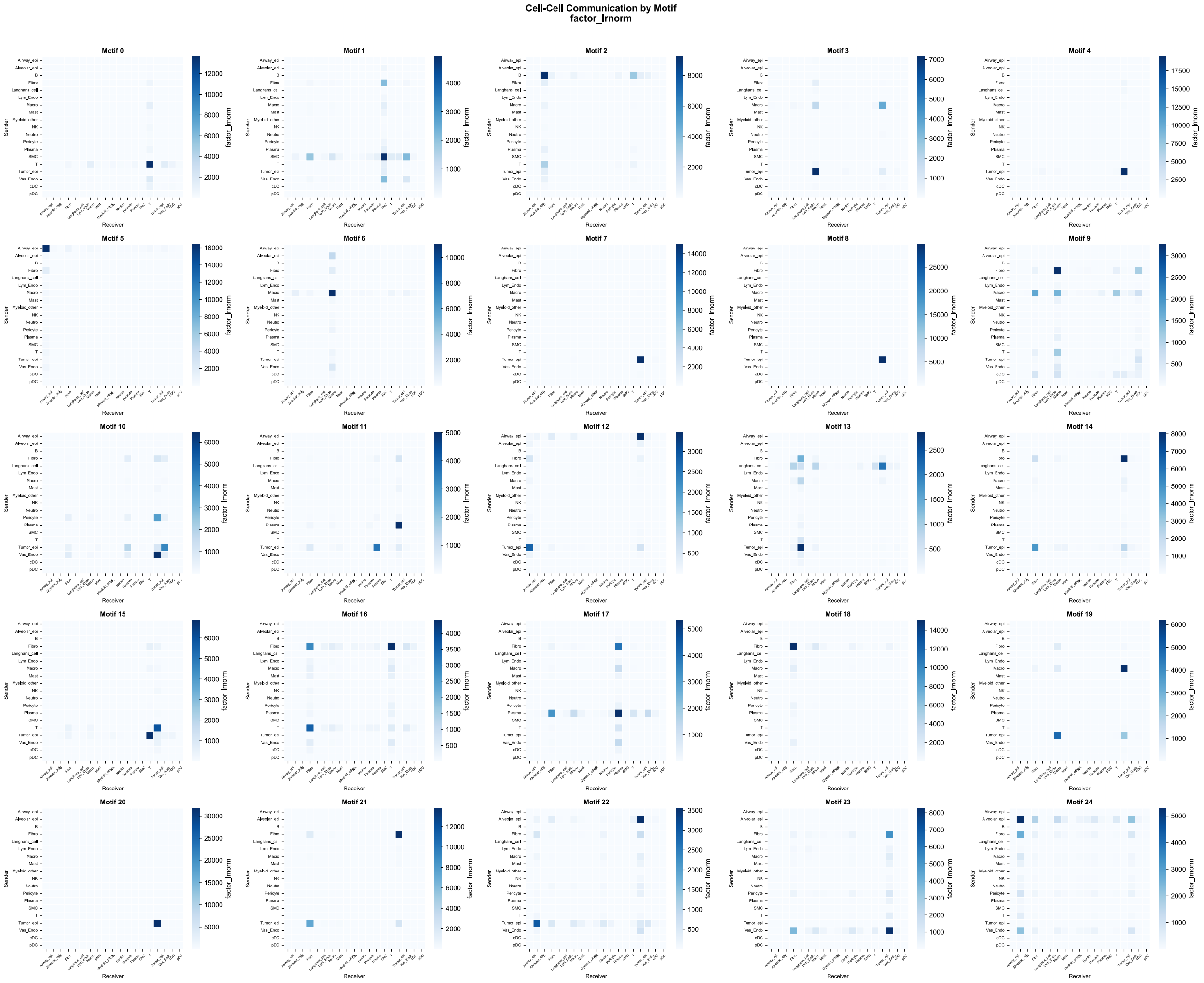


**Supplementary Figure 7: Cell-type-level factor summary across LUAD motifs.** Heatmap showing the aggregated normalized factor value per cell type per motif. For each motif, factor values across all LRIs involving a given cell type (as sender or receiver) were summed. Rows represent cell types and columns represent motifs.

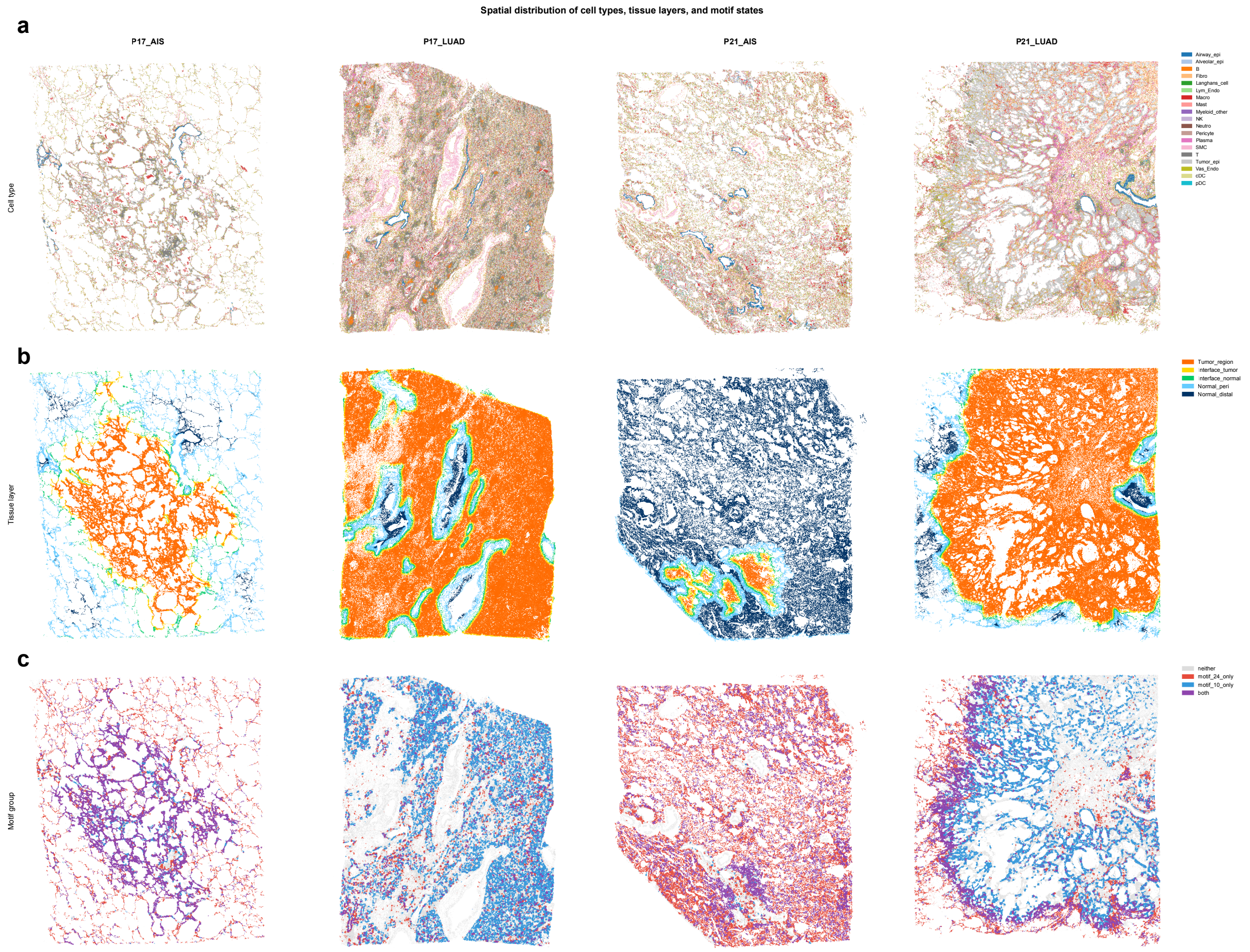


**Supplementary Figure 8: Spatial distribution of motif states and tissue layers.** Spatial scatter plots of all cells across four samples. Top row: cells colored by motif group membership (motif 24 only, red; motif 10 only, blue; both, purple; neither, gray). Bottom row: the same cells colored by tissue layer annotation (tumor region, interface tumor, interface normal, peri-tumoral normal, distal normal). Tissue layers were defined based on pathological annotations from Peng et al. ( ? ). Each column corresponds to one sample.

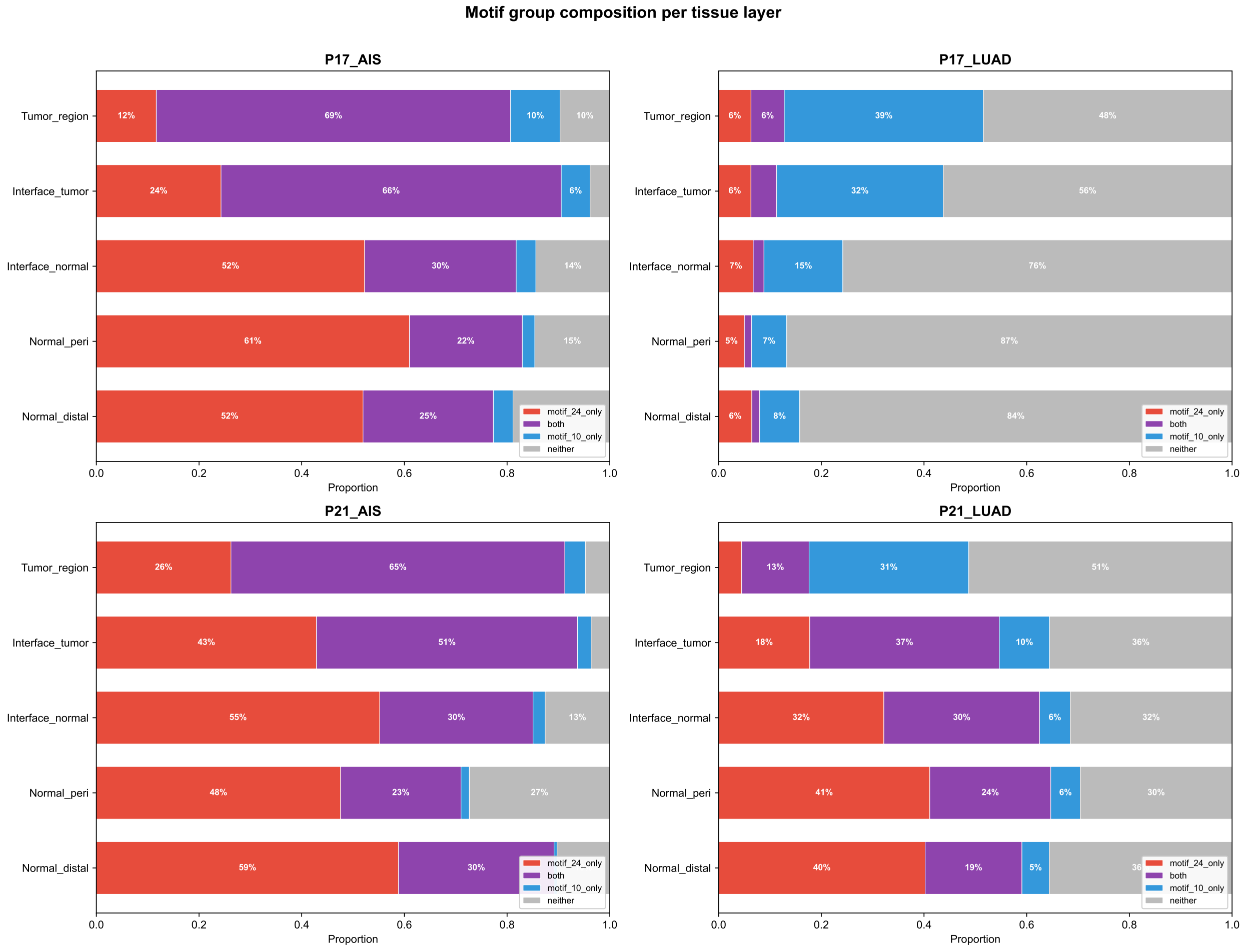


**Supplementary Figure 9: Motif group composition across tissue layers.** Stacked horizontal bar charts showing the proportion of cells assigned to each motif group (motif 24 only, both, motif 10 only, neither) within each tissue layer, stratified by sample. Tissue layers are ordered from tumor core (top) to distal normal (bottom). Percentages are annotated for groups exceeding 5% of the layer.

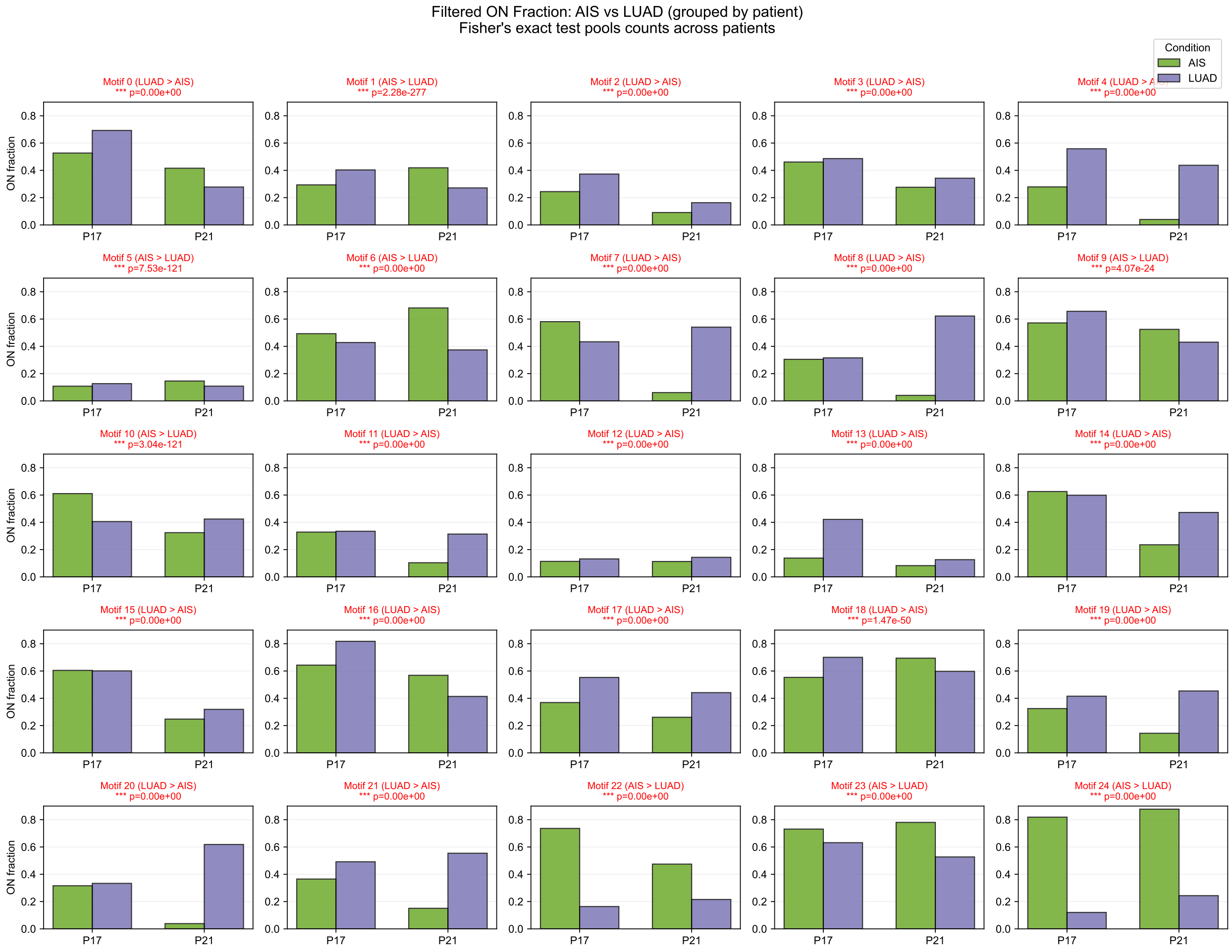


**Supplementary Figure 10: Comparison of motif activation frequencies between AIS and LUAD.** Grouped bar charts showing the ON fraction (proportion of cells in the positive state) for each of 25 motifs, with patients (P17, P21) on the $x$-axis and bars colored by condition (AIS, green; LUAD, purple). Statistical significance was assessed by two-sided Fisher's exact test on cell counts pooled across patients. Asterisks denote significance levels ($*\,p < 0.05$, $**\,p < 0.01$, $*\,*\,*\,p < 0.001$); ns, not significant.

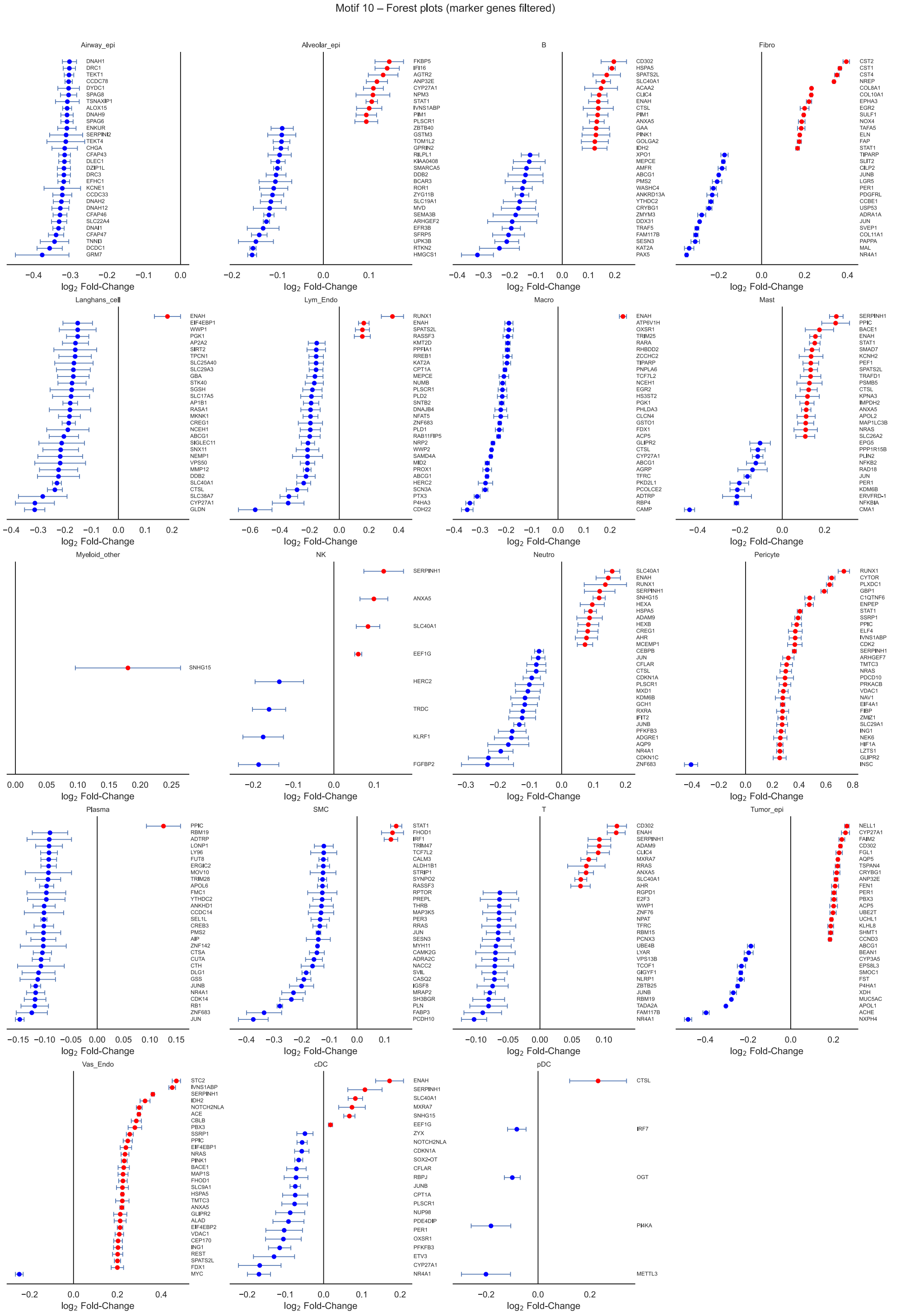

**Supplementary Figure 11: ALARMIST downstream impacts of the tumor vasculature motif (motif 10) across cell types.** Forest plots of significant downstream impact genes associated with motif 10 activity, shown separately for each cell type. Each panel displays the top genes ranked by absolute $\log_2$ fold-change, with red points indicating upregulation and blue points indicating downregulation in motif-active cells. Horizontal bars represent 95% confidence intervals from the ALARMIST Poisson generalized linear model. Ligand and receptor genes used in motif construction are excluded by design; marker genes of other cell types are additionally excluded to avoid confounding from transcript diffusion.

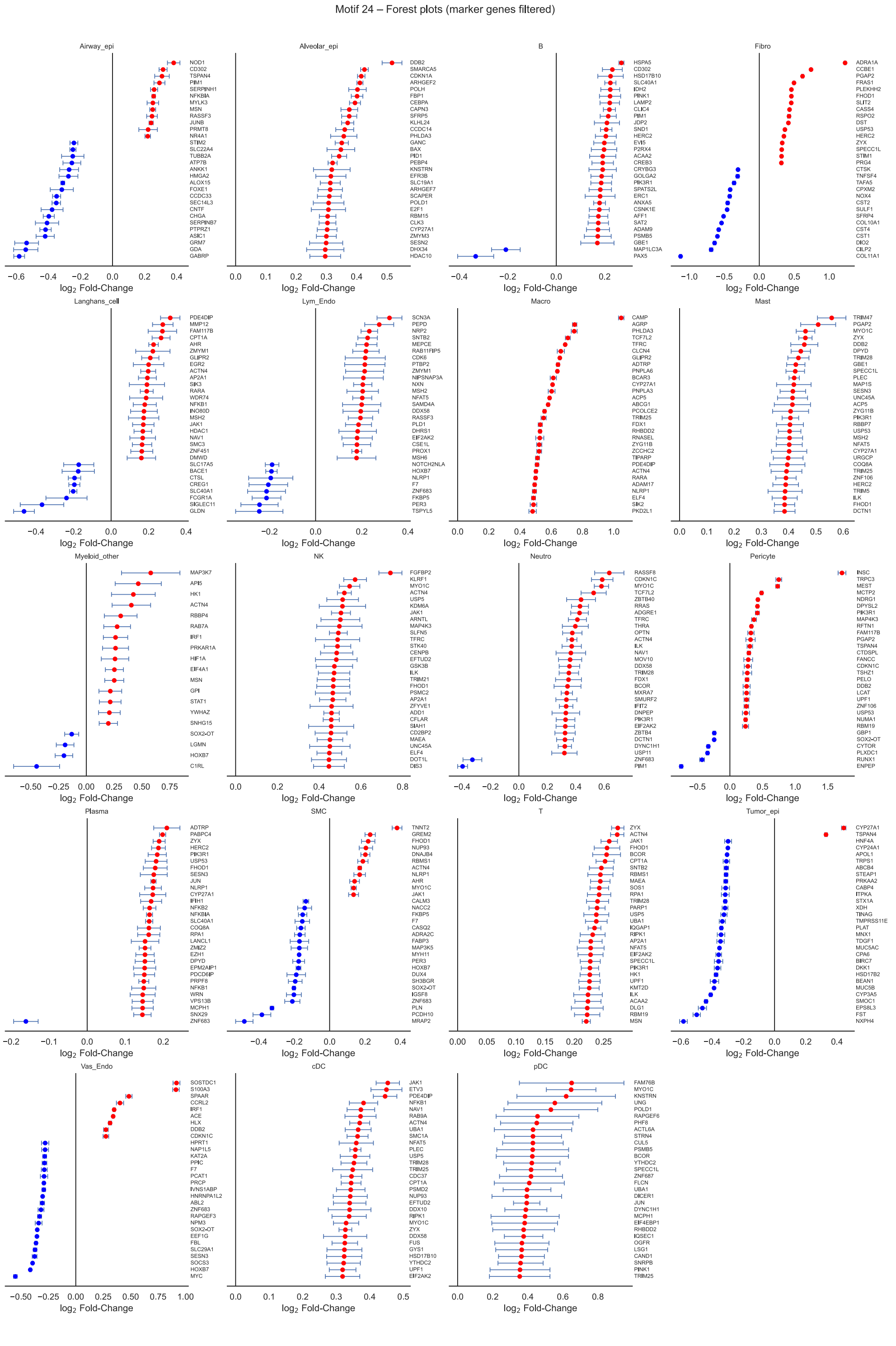


**Supplementary Figure 12: ALARMIST downstream impacts of the healthy vasculature motif (motif 24) across cell types.** Forest plots of significant downstream impact genes associated with motif 24 activity, shown separately for each cell type. Display conventions as in Figure 11.

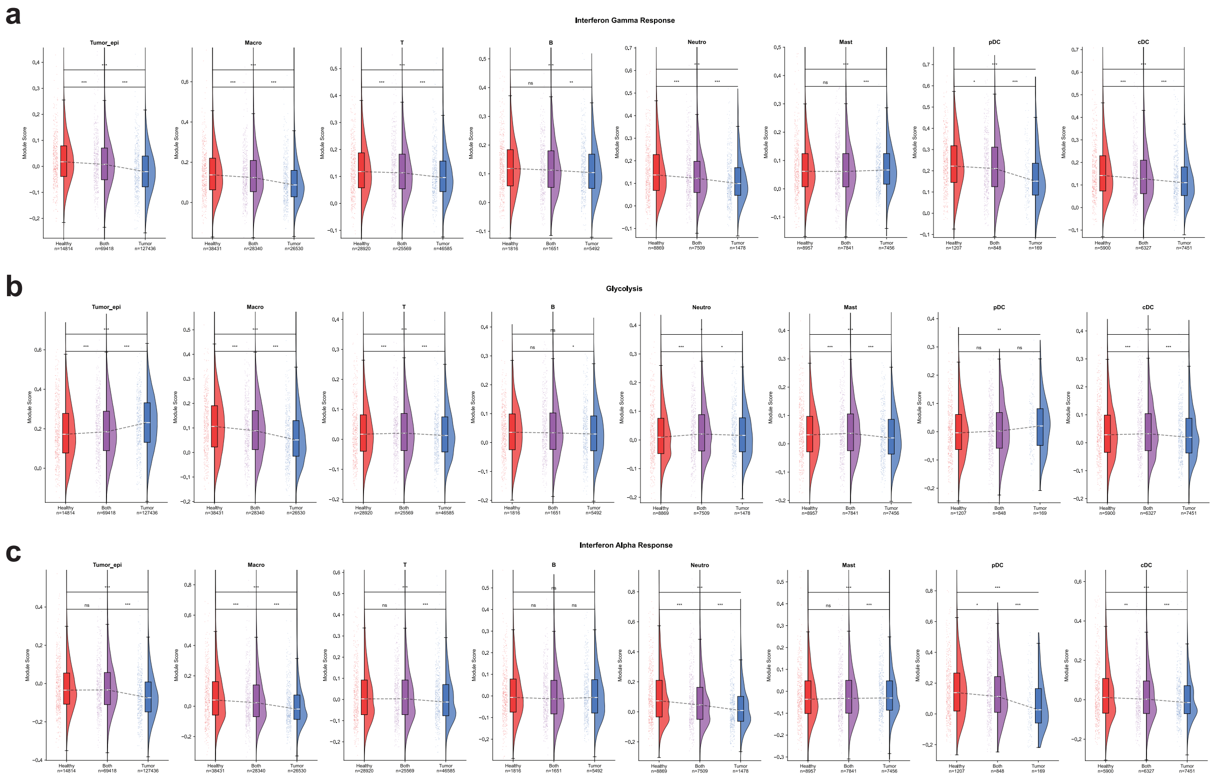

**Supplementary Figure 13: Module scores for key immune and metabolic pathways stratified by vascular motif state across immune cell types and tumor. a.** Interferon-$\gamma$ Response, **b.** Glycolysis, and **c.** Interferon Alpha Response Hallmark module scores for cells classified by vascular motif state (Healthy, healthy vasculature motif ON only; Both, both motifs ON; Tumor, tumor vasculature motif ON only). Each panel shows one cell type; violin plots display the distribution of per-cell module scores, with boxplots indicating median and interquartile range. Sample sizes are shown below each group. Statistical comparisons between adjacent groups were performed using two-sided Wilcoxon rank-sum tests ($***p < 0.001$, $**p < 0.01$, $*p < 0.05$; ns, not significant); brackets spanning all three groups indicate the overall Healthy-versus-Tumor comparison. IFN-$\gamma$ response and glycolysis differences were most pronounced in macrophages, T cells, and cDCs, whereas pDCs showed no significant difference in glycolysis, consistent with their role as upstream IFN signal sources rather than metabolic effectors. Cells negative for both motifs were excluded.

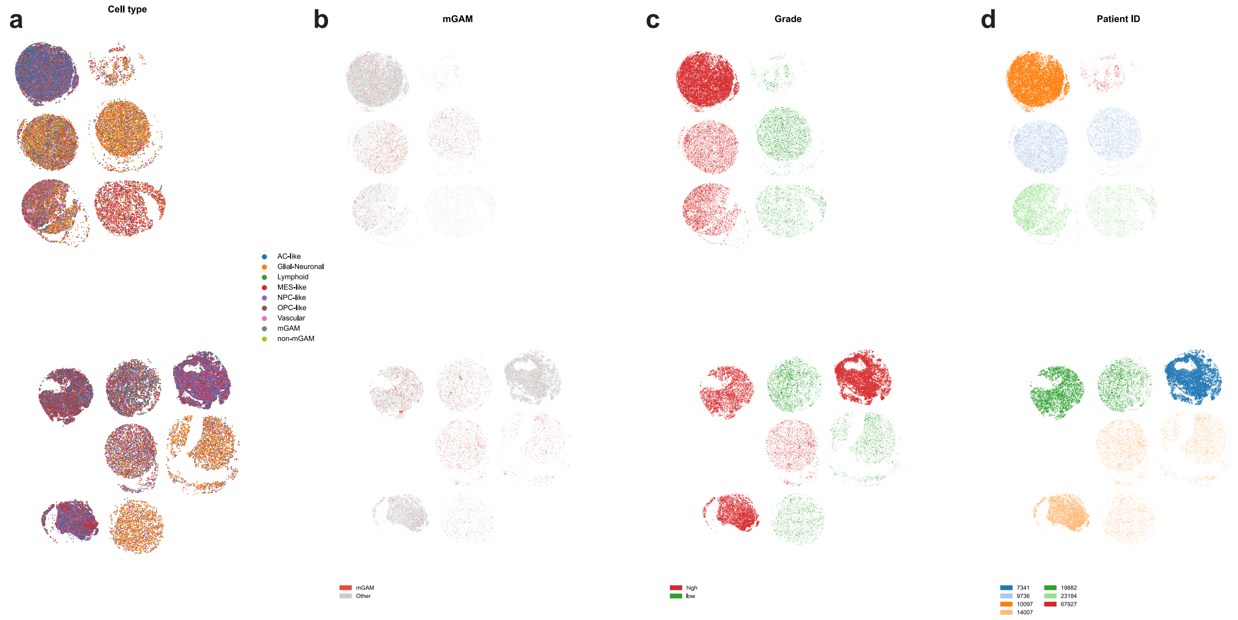


**Supplementary Figure 14: Spatial overview of the GBM tissue microarray.** Spatial maps of all cells across 13 TMA cores. **a.** Cells colored by cell-type annotation. **b.** mGAM cells highlighted in red with all other cell types in gray. **c.** Cells colored by histological grade (high vs. low). **d.** Cells colored by patient ID. Each dot represents a single cell plotted at its spatial centroid coordinate.

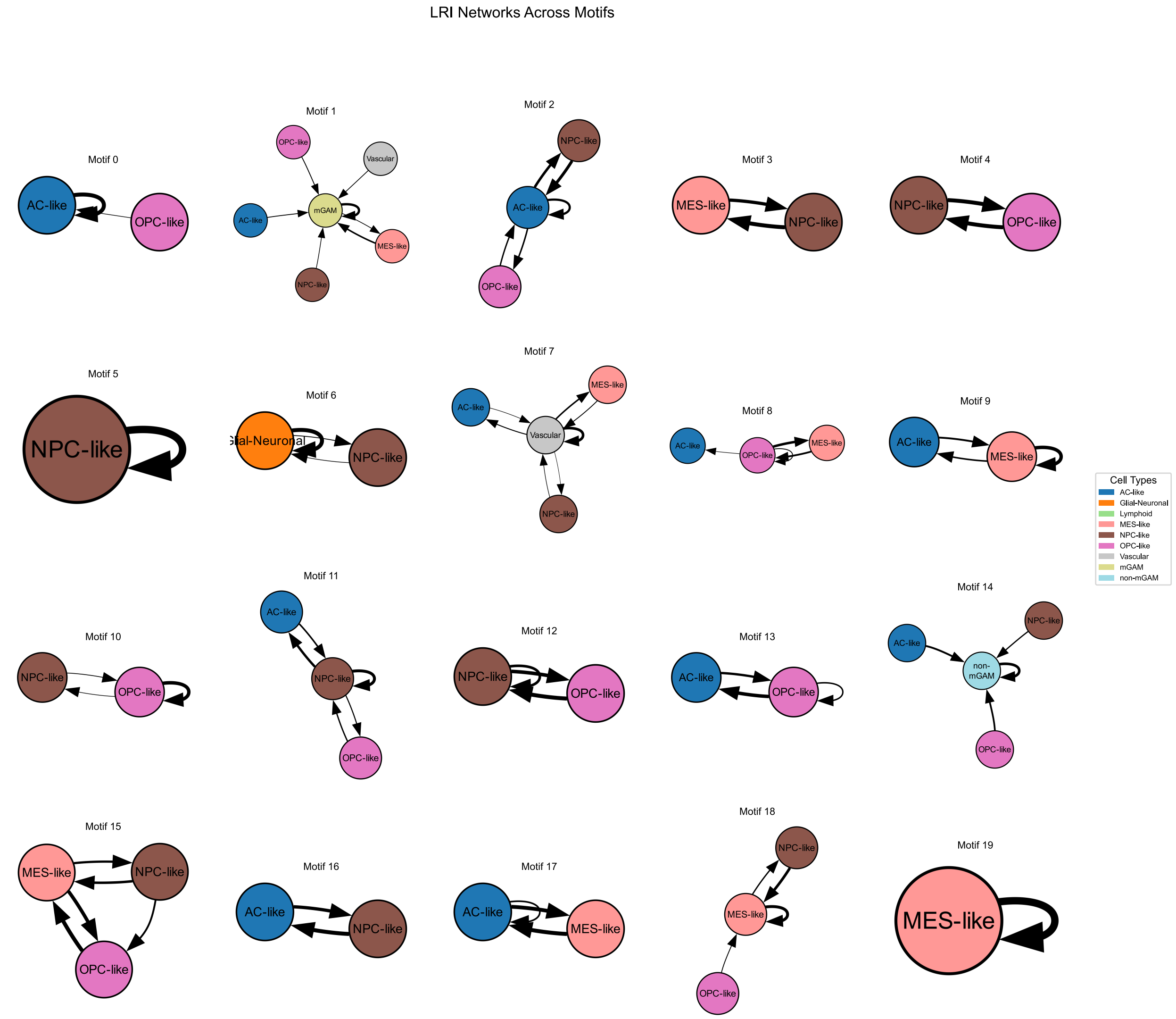


**Supplementary Figure 15: Cell-type interaction networks for all GBM motifs.** Each panel shows one motif as a network graph. For each motif, the top 500 LRIs (ranked by normalized factor value) were aggregated by sender–receiver cell-type pair. Cell types with aggregated factor values exceeding 800 are displayed as nodes, with edges representing LRI interactions between them. Edge thickness reflects the summed factor value.

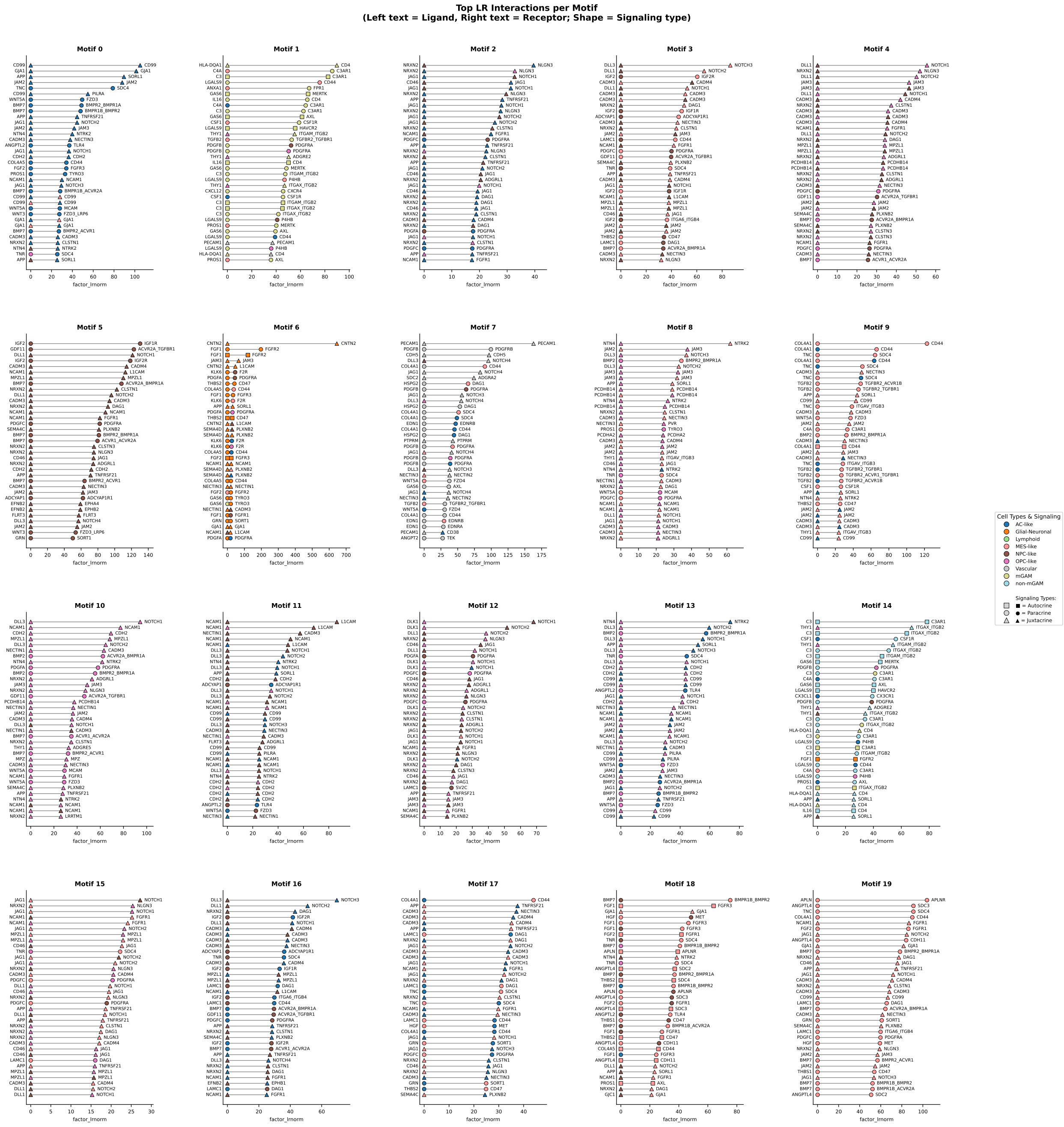


**Supplementary Figure 16: Top 25 LRIs per motif in GBM.** For each motif, the top 25 LRIs ranked by normalized factor value are shown. Each bar represents one LRI, labeled by sender cell type, receiver cell type, and ligand–receptor pair.

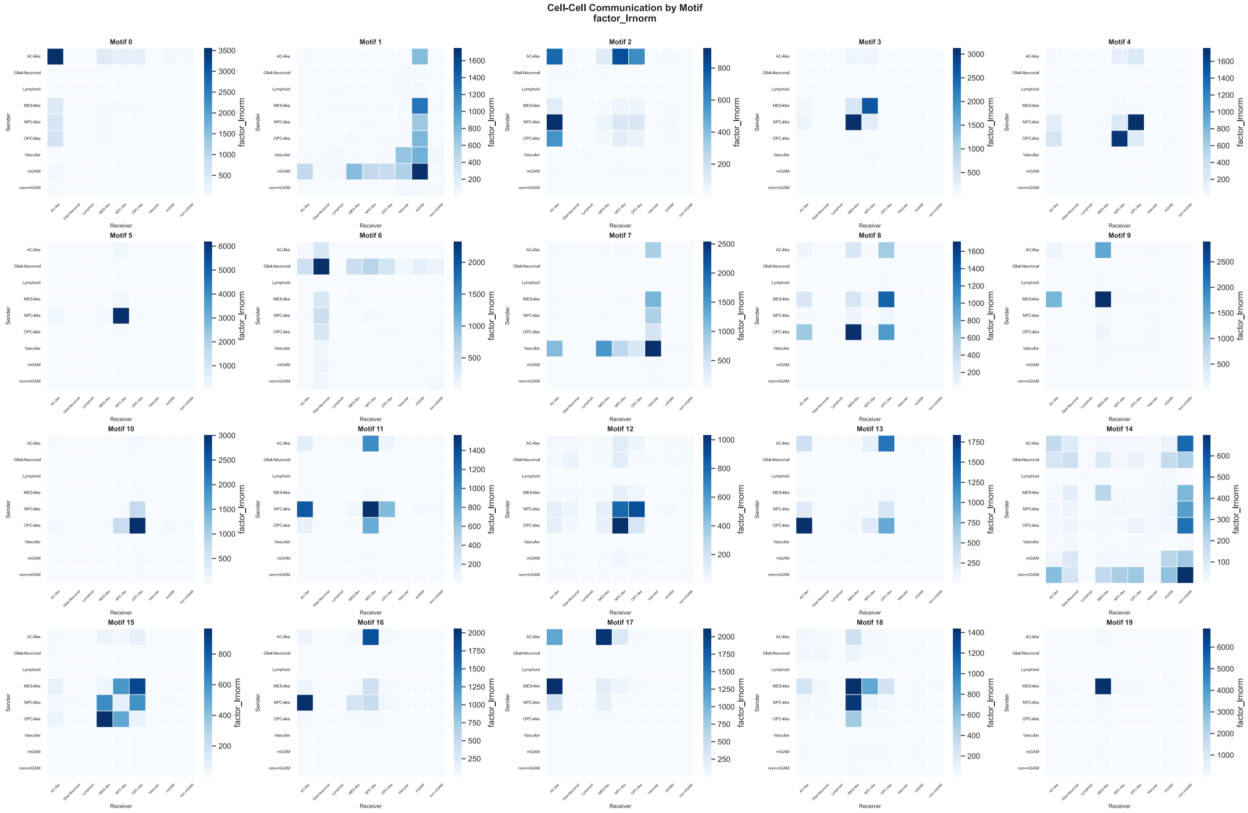


**Supplementary Figure 17: Cell-type-level factor summary across GBM motifs.** Heatmap showing the aggregated normalized factor value per cell type per motif. For each motif, factor values across all LRIs involving a given cell type (as sender or receiver) were summed. Rows represent cell types and columns represent motifs.

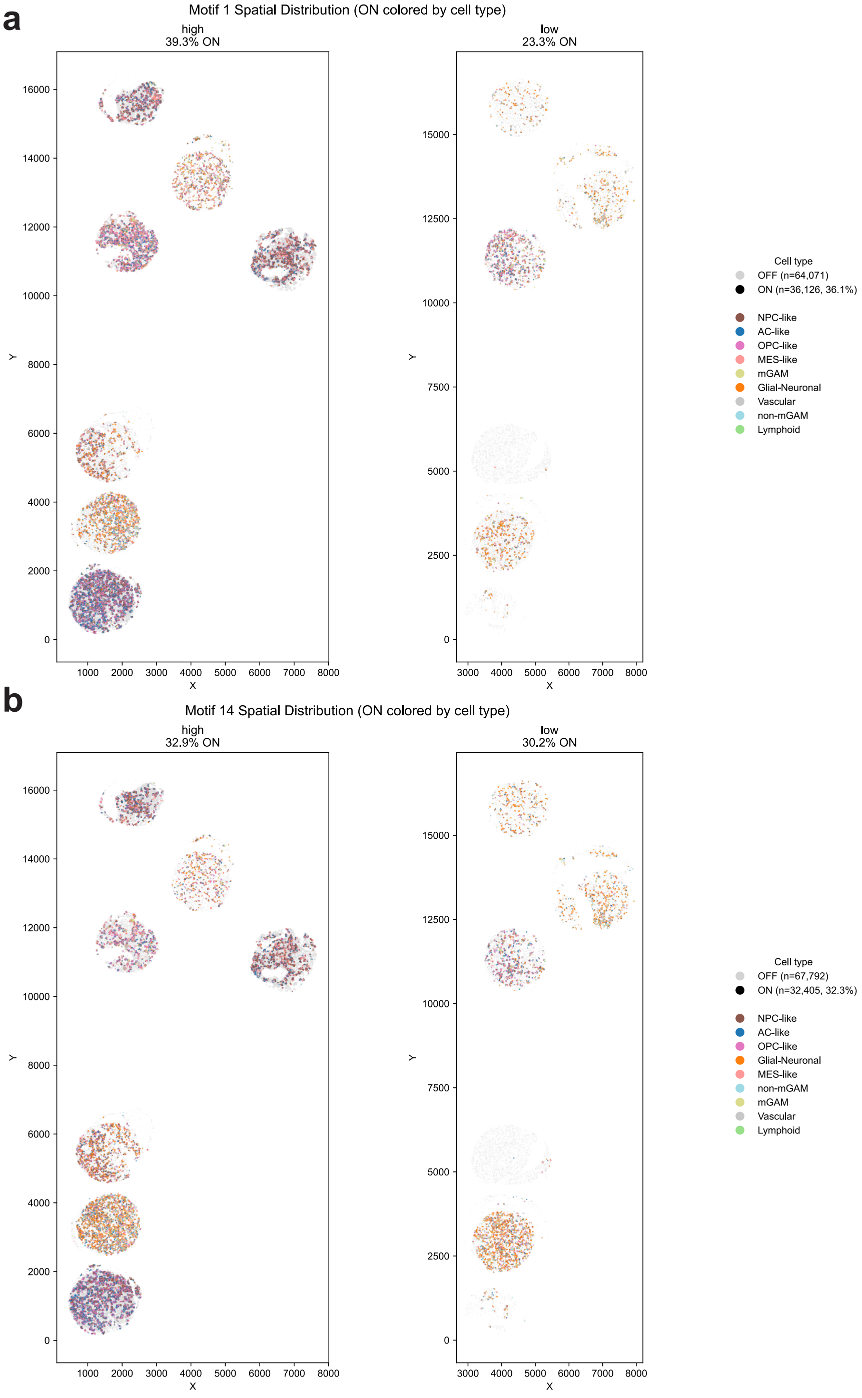


**Supplementary Figure 18: Spatial distribution of motif 1 and motif 14 activation states. a.** Spatial maps of motif 1 ON (positive) and OFF (negative) cells, split by tumor grade (high vs. low). Cells in the ON state are colored by cell type; OFF cells are shown in gray. Motif 1 captures a signaling program enriched for mGAM interactions. **b.** Same layout for motif 14, which captures a distinct intercellular signaling program involving non-MES tumor subtype communication with mGAM.

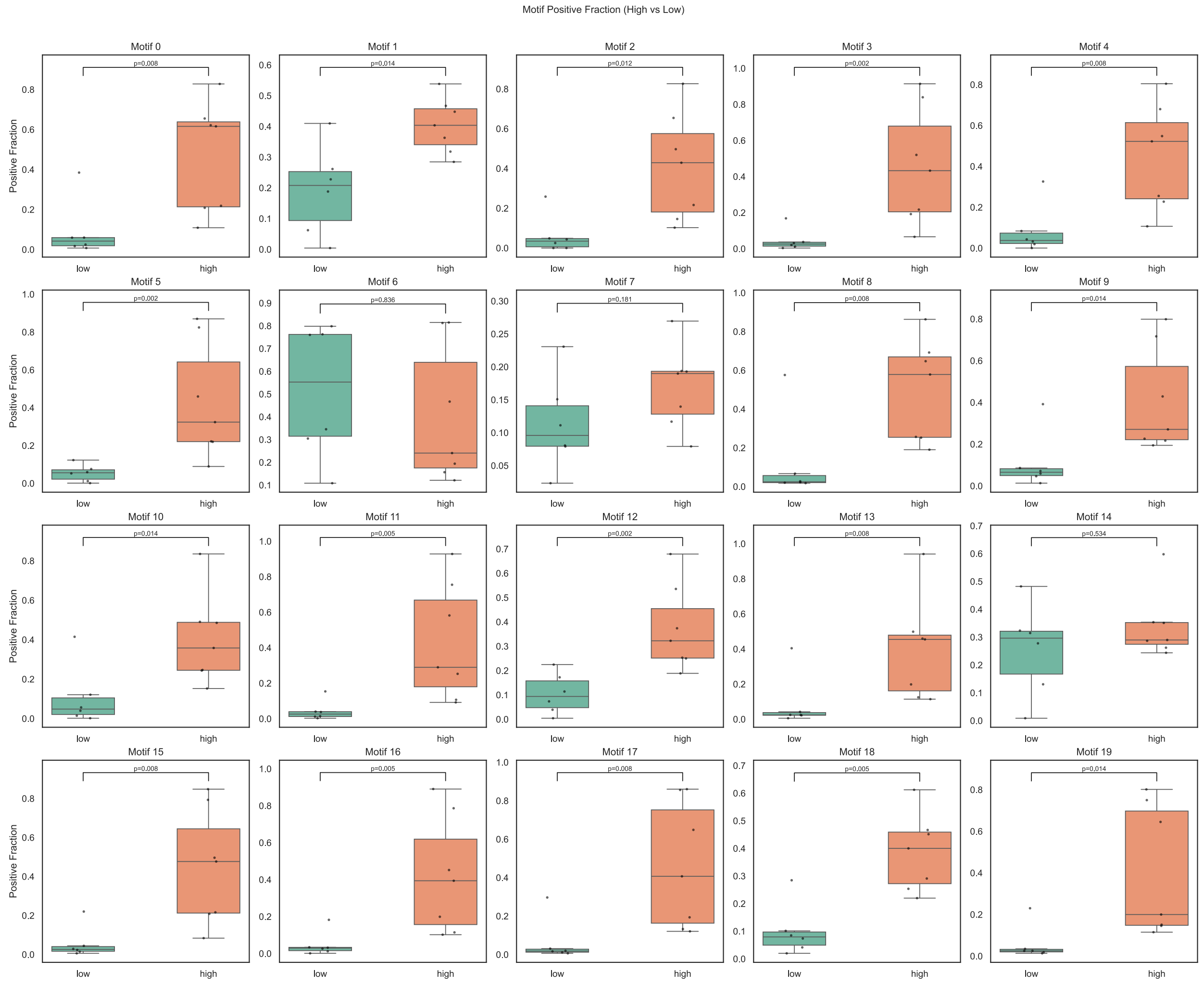


**Supplementary Figure 19: Motif activation frequency by tumor grade across all GBM motifs.** Each panel shows one motif. Boxplots display the fraction of motif-positive cells in low-grade versus high-grade samples. Each dot represents one sample. $P$-values from two-sided Mann–Whitney $U$ tests are annotated above each comparison.

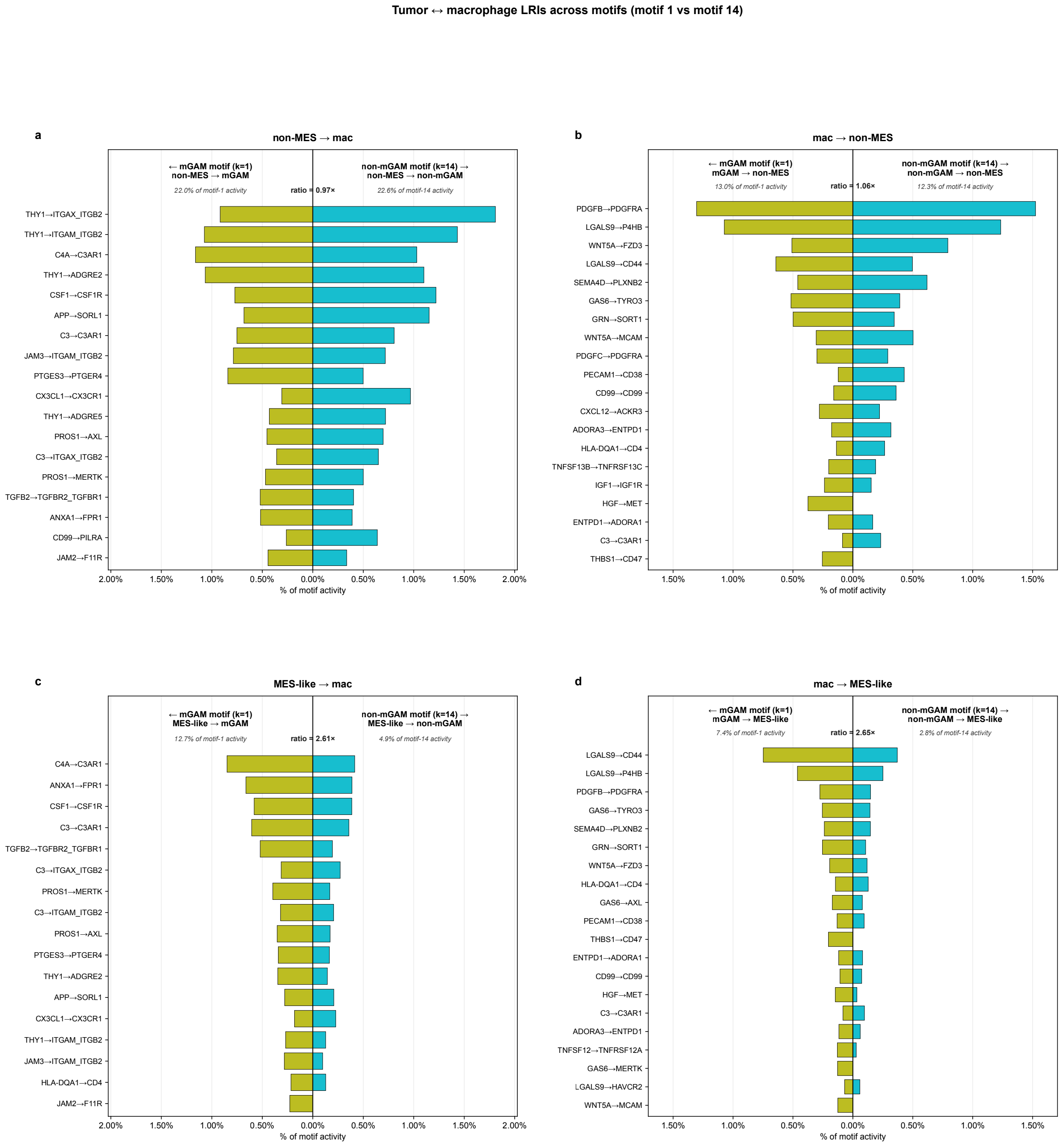


**Supplementary Figure 20: Tumor-macrophage LRI composition is shared on the non-MES axis but diverges on the MES-like axis between the mGAM and non-mGAM motifs.** Butterfly plots comparing the top LRIs on each direction of tumor-macrophage signaling between the mGAM motif (motif 1, left, yellow-green) and the non-mGAM motif (motif 14, right, blue). Panels are split by tumor group (rows: non-MES = AC-like / NPC-like / OPC-like, top; MES-like, bottom) and signaling direction (columns: tumor → macrophage, left; macrophage → tumor, right). Bar length is each LRI's fraction of within-motif total activity, normalized to remove the BPTF factor scale degeneracy across motifs. The header of each panel reports each side's total slice fraction and the motif-1 / motif-14 ratio. Non-MES → macrophage signaling is conserved across motifs (a, b; ratios 0.97× and 1.06×), whereas MES-like ↔ macrophage signaling is approximately 2.6-fold more concentrated in the mGAM motif (c, d; ratios 2.61× and 2.65×).
